# Genome-wide mapping of endogenous topoisomerase cleavage complexes reveals TOP1 at the interface of DNA topological regulation and genome instability

**DOI:** 10.64898/2026.09.30.755763

**Authors:** Shuai Li, Chongyi Chen

## Abstract

Topoisomerase cleavage complexes (TOPcc) are covalent protein–DNA intermediates formed transiently during topoisomerase catalysis and constitute a major class of DNA–protein crosslinks in human cells, yet their endogenous genomic distributions remain difficult to resolve. Here, we developed protein–DNA adduct immunoprecipitation followed by sequencing (pDAIP-seq), a highly sensitive method for detecting endogenous TOPcc that enables their genome-wide profiling under normal physiological conditions without stabilization by topoisomerase poisons. Using pDAIP-seq, we found that camptothecin treatment, required by existing TOP1cc profiling methods, not only increased TOP1cc abundance but also substantially altered its genomic distribution. Endogenous TOP1cc were globally reduced upon transcriptional suppression and increased upon TOP2A depletion, supporting TOP1cc regulation by the maintenance of genome-wide DNA topological homeostasis. Interestingly, both the regional distribution and nucleotide-scale positioning of TOP1cc, including the positions of TOP1 catalytic cleavage sites, were largely specified by local DNA sequence context. By extending pDAIP-seq to capture transient covalent protein–DNA intermediates involved in DNA repair, including TDP1–, PARP1–, and KU70–DNA adducts, we further identified endogenous TOP1cc as a potential major source of spontaneous DNA breaks in human cells. Finally, pDAIP-seq also enabled genome-wide mapping of TOP2cc, TOP3cc, and SPO11–DNA covalent intermediates in mouse testis undergoing programmed meiotic DNA cleavage. Together, these findings establish pDAIP-seq as a broadly applicable method for the mapping of covalent protein–DNA intermediates and reveal endogenous TOP1cc as a key intermediate between DNA topological regulation and human genome instability.

## Main Text

DNA topological homeostasis is continuously challenged by chromatin activities including transcription, replication, chromosome segregation, and genome folding(*1*). To maintain topological homeostasis across the genome, DNA topoisomerases resolve DNA supercoiling tension through the catalytic cycles of DNA cleavage and religation. These reactions generate transient covalent protein–DNA intermediates known as topoisomerase cleavage complexes (TOPcc)(*2–4*). Although topoisomerase catalysis is essential for genome regulation and function, its cleavage intermediates can become potentially mutagenic lesions when DNA religation fails, resulting in abortive catalytic cycles and trapped TOPcc(*2*, *5*, *6*). Emerging evidence supports the contribution of such events to genome instability in diverse biological contexts. TOP1 activity has been linked to a characteristic transcription-associated deletion signature in both cancer and the germline(*7*). Aberrant TOP1-mediated mutagenesis has also recently been identified as a major source of somatic mutations in neurons from patients with amyotrophic lateral sclerosis (ALS), frontotemporal dementia (FTD), and Alzheimer’s disease(*8*). TOP2B cleavage complexes can be trapped during the confined migration of developing neurons, generating double-strand breaks that are normally repaired(*9*). Topoisomerase-linked DNA transactions have further been implicated in replication-associated DNA damage responses and the maintenance of oncogenic extrachromosomal DNA(*10*, *11*). Together, these findings raise the possibility that transient TOPcc generated during topoisomerase catalysis under normal physiology constitute an intrinsic link between DNA topological regulation and genome instability.

Despite extensive biochemical and structural understanding of topoisomerase catalysis and TOPcc formation, the genome-wide landscape of endogenous TOPcc generated as transient catalytic intermediates under normal physiological conditions remains difficult to study. A major challenge is their transient nature, since many existing genomic approaches rely on topoisomerase poisons to stabilize TOPcc before detection, that increase TOPcc abundance but may also alter its genomic distribution, potentially obscuring its endogenous landscape (*12–16*). Consequently, fundamental questions remain unresolved regarding where TOPcc form across the genome, what determines their endogenous abundance and genomic distribution, and to what extent they contribute to genome instability.

Here, we develop protein–DNA adduct immunoprecipitation followed by sequencing (pDAIP-seq), a platform for genome-wide mapping of covalent protein–DNA complexes while preserving their endogenous genomic information without poison-mediated stabilization. Using pDAIP-seq, we map endogenous TOP1cc across the human genome and identify distinct principles governing their abundance and distribution. We find that TOP1cc levels are regulated by genome-wide DNA topological stress, whereas their regional distribution and nucleotide-scale positioning are primarily determined by local DNA sequence context. We further extend pDAIP-seq to map covalent protein–DNA intermediates associated with DNA repair and identify TOP1cc as a potential major endogenous source of spontaneous DNA damage in human cells. Our findings establish a framework for studying transient protein–DNA intermediates under normal physiological conditions and position TOP1cc at the interface between DNA topological homeostatic maintenance and genome instability.

### pDAIP-seq maps endogenous TOP1 cleavage complexes (TOP1cc) without poison stabilization

TOP1 resolves torsional stress through a reversible catalytic cycle in which the enzyme cleaves one DNA strand, becomes covalently linked to the 3’ DNA end, permits controlled rotation, and then religates the strand (Fig. 1A). The short lifetime of this transient intermediate makes TOP1cc difficult to detect without topoisomerase poison stabilization in existing assays. To overcome this challenge, we developed protein–DNA adduct immunoprecipitation followed by sequencing (pDAIP-seq) to recover covalent complexes without drug-mediated stabilization.

**Fig. 1.**
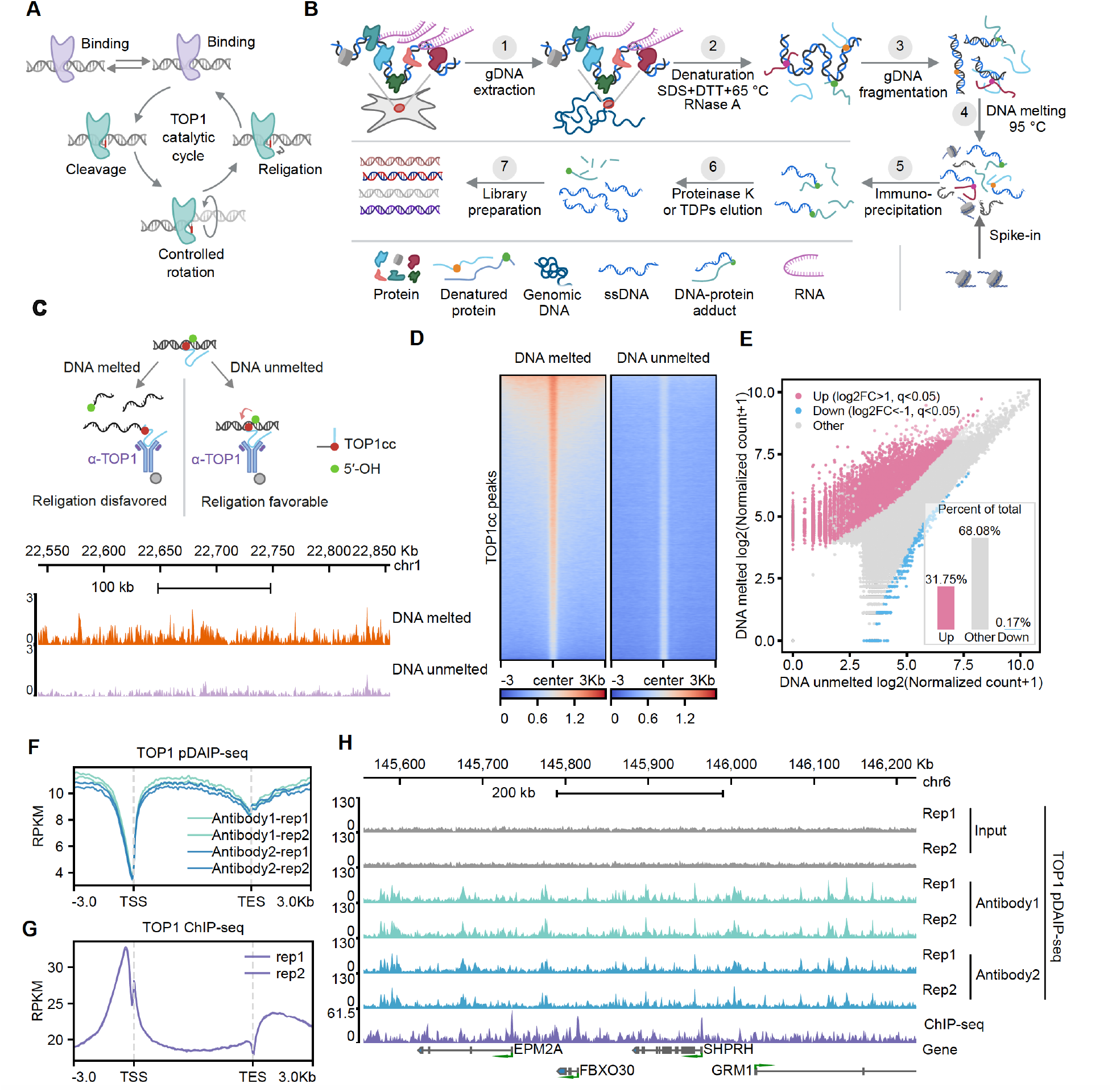
pDAIP-seq maps endogenous TOP1 cleavage complexes without exogenous poison stabilization. (**A**) Catalytic cycle of human TOP1, including DNA binding, single-strand cleavage and covalent TOP1-DNA intermediate formation, controlled rotation, and religation. (**B**) Schematic workflow of protein–DNA adduct immunoprecipitation followed by sequencing (pDAIP-seq). (**C**) Schematic and representative region on chromosome 1 comparing TOP1 pDAIP-seq signal with or without DNA melting before immunoprecipitation. DNA melting disrupts the duplex geometry favorable for religation and is expected to reduce post-isolation reversal of TOP1 cleavage complexes. (**D**) TOP1cc-centered heatmaps comparing DNA-melted and DNA-unmelted preparations across 3-kb flanking regions. (**E**) Differential recovery of TOP1cc peaks between DNA-melted and DNA-unmelted preparations. Pink and blue points indicate regions with log2 fold change > 1 or < −1, respectively, at q < 0.05; the inset shows the percentage of regions in each class. (**F**) Metagene profiles of TOP1 pDAIP-seq signal across gene bodies and 3-kb flanking regions for two biological replicates generated with each of two TOP1 antibodies. (**G**) Metagene profiles of TOP1 ChIP-seq signal across the same gene-centered interval for two biological replicates. (**H**) Representative region on chromosome 6 showing input, TOP1 pDAIP-seq replicates generated with the two antibodies, TOP1 ChIP-seq, and gene annotations. TSS: transcription start site; TES: transcription end sites: PRKM: reads per kilobase per million mapped reads.

Genomic DNA was extracted under protein-denaturing conditions, treated with RNase A, fragmented, heated to melt duplex DNA, and immunoprecipitated with a TOP1 antibody in the presence of a spike-in reference. Adduct-bearing DNA was released with proteinase K or tyrosyl-DNA phosphodiesterase before library preparation (Fig. 1B). Assay specificity was supported by IgG immunoprecipitation and proteinase K pretreatment controls, both of which reduced TOP1cc recovery (fig. S1, A and B). As an independent validation of catalytic specificity, we performed GFP-directed pDAIP-seq in cells expressing doxycycline-inducible

GFP-TOP1 or the catalytically inactive GFP-TOP1^Y723F^ mutant. Although both proteins were expressed, only GFP-TOP1 showed pDAIP-seq recovery but not GFP-TOP1^Y723F^, further supporting selective detection of covalent complex of TOP1cc rather than TOP1 binding (fig. S1, C to E).

Importantly, we noted that because IP requires SDS dilution into Triton X-100, residual TOP1 activity may become less inhibited(*17*). Under these potentially permissive conditions, religation could resume, a reaction that requires duplex DNA contacts and alignment of the free 5′ hydroxyl with the TOP1-linked 3′ DNA end(*18*, *19*). We therefore tested whether DNA melting before immunoprecipitation altered TOP1cc recovery(*18*). DNA melting is expected to disrupt this geometry and reduce the opportunity for reversal, increasing the TOP1cc IP efficiency.

Relative to the DNA-unmelted preparation, the DNA-melted preparation produced stronger signal across representative loci and at TOP1cc peaks genome-wide (Fig. 1, C and D), as well as the pull-down DNA amount (fig. S1B). Differential analysis showed significantly increased recovery at 32% of TOP1cc peaks in the DNA-melted condition (Fig. 1E). Thus, DNA melting indeed increases TOP1cc recovery. This result is consistent with TOP1cc loss through religation during immunoprecipitation, although the experiment does not directly measure TOP1cc religation kinetics.

Using these conditions, reproducible profiles around genes were generated (Fig. 1F), with highly correlated results from both biological replicates and replicates using different TOP1 antibodies (fig. S1F-H). Data-quality fingerprint plots further showed enrichment relative to input for both antibodies (fig. S1I). TOP1cc signal extended across transcription units but decreased sharply at transcription start sites and more modestly near transcription end sites (Fig. 1F). This pattern differed from TOP1 ChIP-seq, which showed prominent TOP1 binding at promoters (Fig. 1G). Concordant signal from the two antibodies was also evident across individual loci of TOP1cc, and different from TOP1 binding sites (Fig. 1H). Peak-length and genome-wide annotation analyses further showed that TOP1cc and total TOP1 peak sets differed in their genomic distributions and length profiles (fig. S1J-L). Together, these results distinguish the genomic distribution of TOP1cc from that of total chromatin-bound TOP1, and establish pDAIP-seq as a reproducible approach for mapping endogenous TOP1cc without poison stabilization for enrichment.

### Camptothecin enriches TOP1cc and alters its endogenous genomic landscape

Having established the endogenous TOP1cc readout, we next asked a question how poison stabilization impact TOP1cc and chromatin. Camptothecin (CPT) stabilizes TOP1cc by inhibiting religation and enriches TOP1cc for detection(*20*). We compared untreated and CPT-treated cells by TOP1 ChIP-seq, TOP1 pDAIP-seq, KAS-seq for single-stranded DNA, and DRIP-Rx, a spike-in-normalized S9.6 assay for R-loops incorporating RNase T1 and RNase III pretreatment to reduce nonhybrid RNA background (Fig. 2A). We found that CPT treatment indeed produced a pronounced increase in TOP1cc signal at representative loci and across transcription units (Fig. 2, B and C), leading to a genome-wide TOP1cc distribution increase by a median of 16-fold (fig. S2B).

**Fig. 2.**
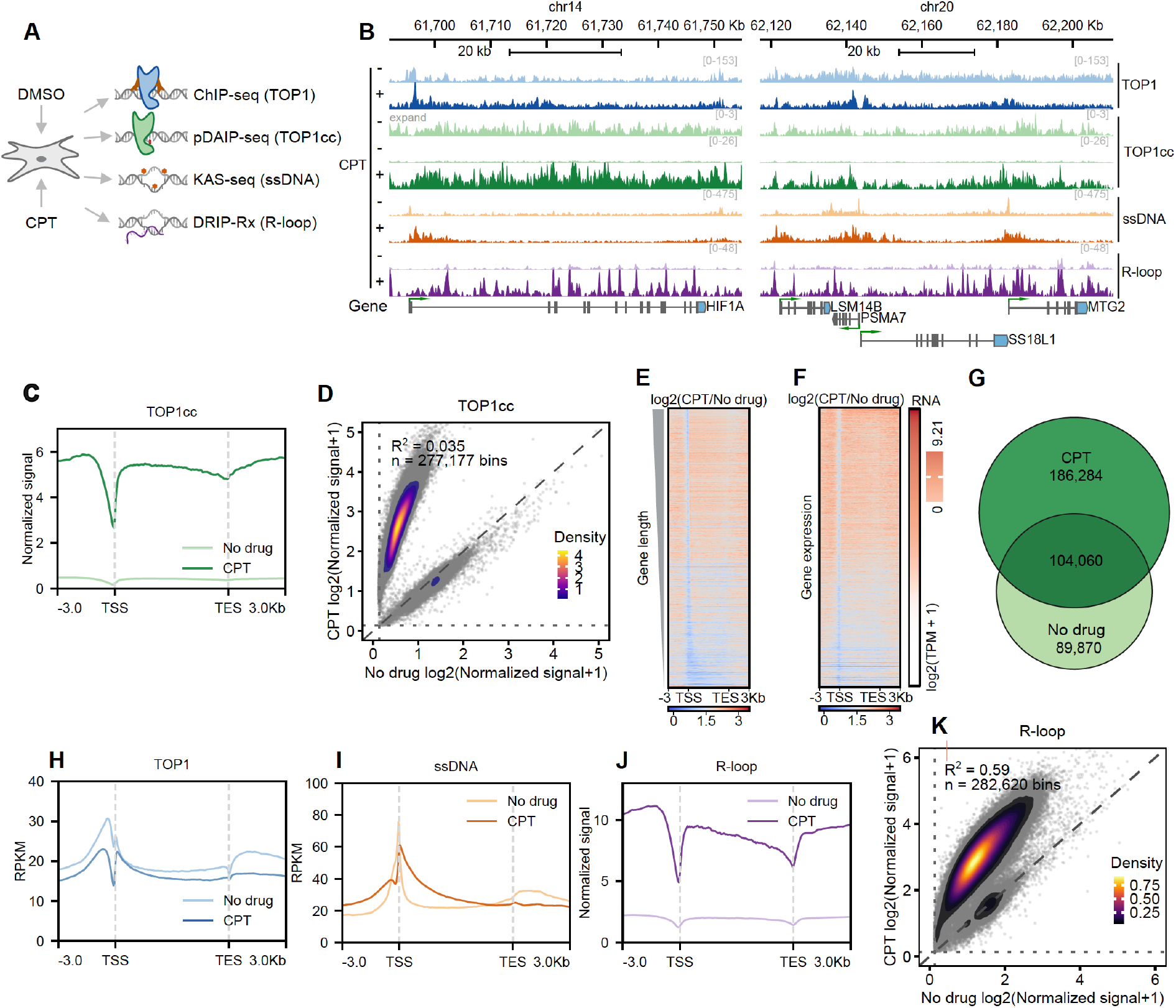
Camptothecin enriches TOP1cc for capture but remodels its endogenous genomic landscape. **(A)** Experimental design comparing no-drug (DMSO) and camptothecin (CPT)-treated cells by TOP1 ChIP-seq, TOP1 pDAIP-seq, KAS-seq for single-stranded DNA (ssDNA), and DRIP-Rx for R-loops. **(B)** Representative regions on chromosomes 14 and 20 showing TOP1, TOP1cc, ssDNA, R-loop, and gene tracks under no-drug and CPT conditions. **(C)** Metagene profiles of TOP1cc signal across transcription units and 3-kb flanking regions under no-drug and CPT conditions. **(D)** Genome-wide comparison of TOP1cc signal between no-drug and CPT conditions across 277,177 genomic bins. **(E)** Gene-centered heatmap of the change in TOP1cc signal following CPT treatment, shown as log2(CPT/no drug), across transcription units and 3-kb flanking regions, with genes arranged by gene length. **(F)** Gene-centered heatmap of TOP1cc log2(CPT/no drug) signal with genes arranged by gene expression; corresponding RNA expression [log2(TPM + 1)] is shown alongside. **(G)** Area-proportional overlap of TOP1cc peaks between conditions, showing 186,284 CPT-specific, 104,060 shared, and 89,870 no-drug-specific regions. **(H-J)** Metagene profiles of total TOP1 occupancy **(H)**, ssDNA **(I)**, and R-loop signal **(J)** across transcription units and 3-kb flanking regions under no-drug and CPT conditions. **(K)** Genome-wide comparison of R-loop signal between no-drug and CPT conditions across 282,620 genomic bins. TSS, transcription start site; TES, transcription end site; RPKM, reads per kilobase per million mapped reads; TPM, transcripts per million.

Importantly, CPT did more than TOP1cc enrichment to elevate its global level for detection. Signals across genomic bins were poorly correlated between CPT treated and untreated conditions (Fig. 2D). Gene-centered heatmaps further showed broad CPT-induced changes differentially affecting genes with different gene lengths or expression levels (Fig. 2, E and F). Peak analysis identified many regions shared between conditions, together with even more peaks that are CPT-specific or untreated-specific (Fig. 2G). CPT also changed total TOP1 occupancy and increased the gene-associated single-stranded DNA and R-loop signals (Fig. 2, H to J), further indicating fundamental alterations of the chromatin including the TOP1cc genomic distribution. R-loop distribution was also different between conditions (Fig. 2K), and RNase H treatment strongly reduced the DRIP-Rx signal, supporting its assignment to RNA-DNA hybrids (fig. S2, C to F). Overall, these results are consistent with classical biochemical work and recent imaging results showing that CPT alters TOP1cc formation sequences and nuclear TOP1 dynamics(*14*, *21*).

Clustering TOP1cc peaks by their response to CPT revealed a graded increase across response groups and linked the altered TOP1cc map to changes in TOP1 occupancy, single-stranded DNA, and R-loops (fig. S2, G to J). These effects further indicate that poison does not simply amplify the untreated TOP1cc population but generates a pharmacologically remodeled landscape, demonstrating the need for drug-free mapping to investigate TOPcc.

### TOP1cc is globally regulated by the maintenance of genome-wide DNA topological homeostasis

Next, we asked how endogenous TOP1cc is regulated under normal physiological conditions. An obvious possibility is that TOP1cc simply accumulates wherever TOP1 is recruited. However, some biochemical evidence of TOP1 and TOP1cc argued against this model. Negative DNA supercoiling can promote local strand separation and G-quadruplex formation at sequence-permissive loci. Human TOP1 can promote G-quadruplex formation from G-rich single-stranded DNA and bind to G-quadruplexes, whereas G-rich single-stranded and G-quadruplex-forming substrates preferentially bind TOP1 but inhibit its cleavage of the DNA (*22*, *23*). and with evidence that chromosomal G-quadruplexes can retain TOP1 while reducing local relaxation activity(*24*).

To test this possibility, we compared our endogenous TOP1cc with total TOP1 occupancy, DNA supercoiling, single-stranded DNA, and G-quadruplex signal. TOP1 occupancy correlated strongly with both negative supercoiling and G-quadruplex signal, whereas TOP1cc showed essentially no genome-wide correlation with total TOP1 or G-quadruplex signal, and a weak inverse association with the measured supercoiling signal (fig. S3, A to F). Gene-centered profiles, representative loci, and supercoiling-centered heatmaps further showed that TOP1cc distribution differs from that of TOP1 binding, single-stranded DNA, and G-quadruplex signal (fig. S3, G to K). The strong association of G-quadruplex signal with TOP1 binding but not with TOP1cc is consistent with TOP1 recruitment or retention in structural contexts that do not proportionately accumulate TOP1cc. Thus, TOP1 recruitment does not directly predict the accumulation of TOP1cc.

We therefore turned from where TOP1 binds to the topological demand for its activity. Specifically, we examined the relationship between TOP1cc level and genome-wide DNA supercoiling tension, which topoisomerase catalysis continuously resolve in order to maintain the DNA topological homeostasis across the human genome.

Because transcription is a major contributor to genome-wide DNA supercoiling (*25*, *26*), we used acute transcription perturbations to test whether changes in DNA topological stress regulate TOP1cc abundance. NAA treatment efficiently depleted degron-tagged POLR2A (Fig. 3, A and B), and POLR2A depletion caused a broad genome-wide reduction in TOP1cc (Fig. 3C and D, fig. S4A). Differential analysis identified approximately 282,000 regions with decreased TOP1cc and only 155 with increased signal, and peak-centered heatmaps showed the same broad loss (fig. S4, B and C). The reduction was observed across genes spanning high, medium, low, and silent expression classes (Fig. 3E), indicating that POLR2A depletion primarily reduced the overall TOP1cc abundance across the genome rather than producing a strong expression-dependent redistribution due to local RNAP-TOP1 interactions.

**Fig. 3.**
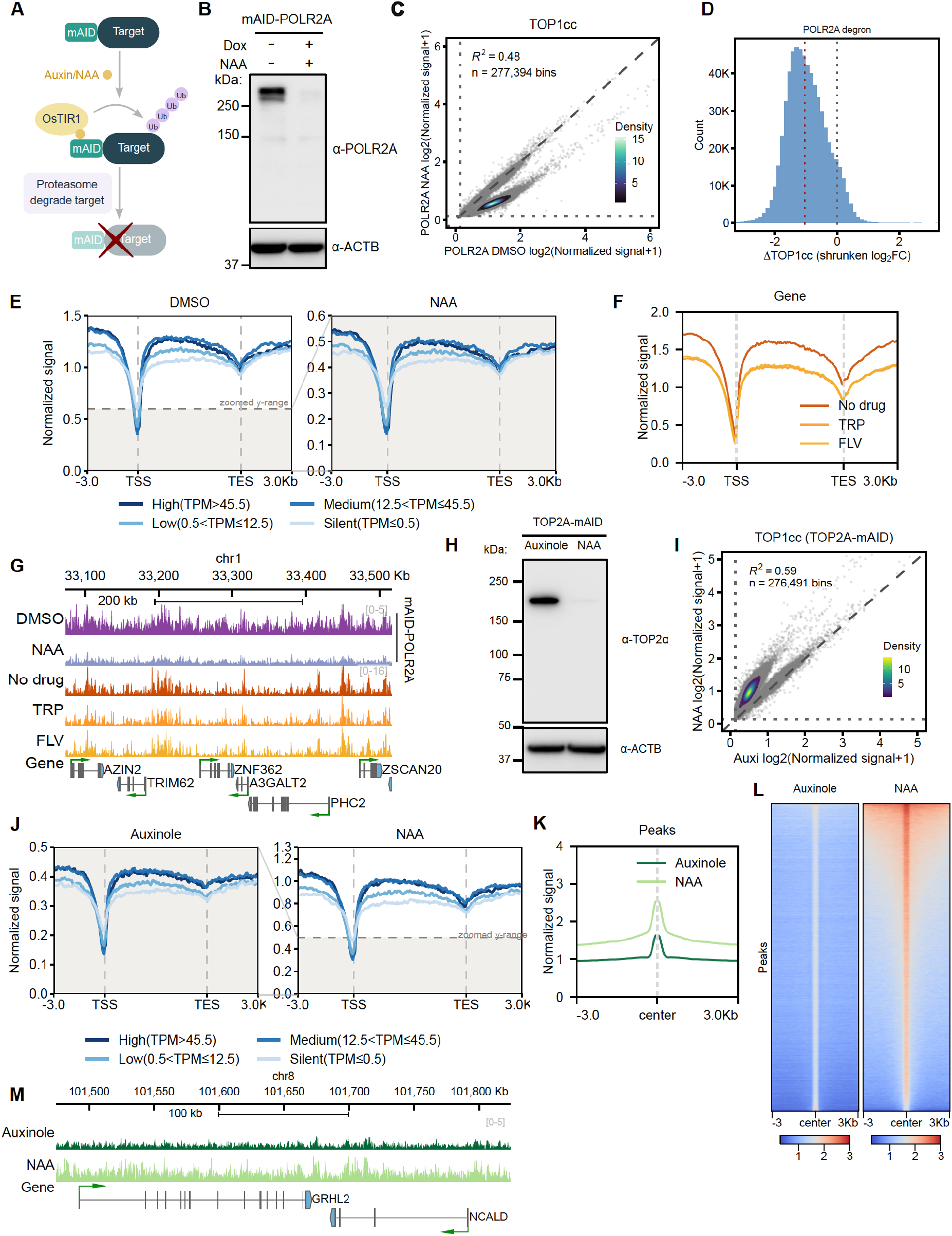
Transcription- and TOP2A-dependent genome-wide DNA topological homeostasis regulates global TOP1cc level. **(A)** Schematic of mini-auxin-inducible degron (mAID)-mediated protein depletion. **(B)** Immunoblot of mAID-POLR2A following the indicated doxycycline (Dox) and 1-naphthaleneacetic acid (NAA) treatments. ACTB serves as a loading control. **(C)** Genome-wide comparison of TOP1cc signal across 277,394 bins before and after POLR2A depletion. **(D)** Distribution of shrunken TOP1cc log2 fold changes following POLR2A depletion. **(E)** Gene-centered TOP1cc profiles for genes stratified by expression level under DMSO and NAA conditions. Expression classes are high (TPM > 45.5), medium (12.5 < TPM ≤ 45.5), low (0.5 < TPM ≤ 12.5), and silent (TPM ≤ 0.5). **(F)** Gene-centered TOP1cc profiles following treatment with triptolide (TRP), flavopiridol (FLV), or no drug. **(G)** Representative chromosome 1 region showing TOP1cc signal following POLR2A depletion or drug inhibition of transcription. **(H)** Immunoblot of mAID-TOP2A following Auxinole or NAA treatment. ACTB serves as a loading control. **(I)** Genome-wide comparison of TOP1cc signal across 276,491 bins before and after TOP2A depletion. **(J)** Gene-centered TOP1cc profiles across the four expression classes under Auxinole and NAA conditions. **(K, L)** Average profiles **(K)** and heatmaps **(L)** centered on pre-existing TOP1cc peaks, comparing Auxinole and NAA conditions across ±3-kb regions. **(M)** Representative chromosome 8 region showing increased TOP1cc signal following TOP2A depletion. TSS, transcription start site; TES, transcription end site; TPM, transcripts per million.

To further determine whether this response was specific to POLR2A depletion, we independently inhibited transcription initiation with triptolide (TRP) or elongation with flavopiridol (FLV). Both treatments reduced TOP1cc across genes with various expression classes including silent ones without any transcription activity (Fig. 3F and fig. S4D), and differential analyses likewise showed predominantly decreased signal (fig. S4E). The TRP- and FLV-induced changes were correlated across genomic bins, whereas either inhibitor showed some bin-level correlation with the POLR2A-degron response (fig. S4, F to H). Representative loci showed the same overall decrease after POLR2A depletion and transcription inhibition (Fig. 3G). Thus, distinct modes of transcriptional suppression converge on reduced global TOP1cc abundance across the genome.

Moreover, if global TOP1cc abundance responds to the demand of supercoiling relaxation for the maintenance of genome-wide DNA topological homeostasis, perturbing another type of topoisomerase should produce a distinct effect. Acute TOP2A depletion elevates the genome-wide DNA supercoiling level(*27*). Indeed, we observed increased TOP1cc genome-wide upon TOP2A depletion. Specifically, with immunoblotting confirming acute depletion of degron-tagged TOP2A (Fig. 3H), we found that TOP1cc were increased genome-wide (Fig. 3I), across transcription units spanning all expression classes including silent ones (Fig. 3J) and around pre-existing TOP1cc peaks (Fig. 3, K and L). Increased TOP1cc was also evident at representative genomic regions (Fig. 3M). The opposite responses to transcriptional suppression and TOP2A depletion support a model in which genome-wide DNA topological homeostasis regulates the global abundance of TOP1cc. However, these perturbations primarily altered the overall TOP1cc abundance, leaving open what determines the local genomic positions at which TOP1cc preferentially forms.

### TOP1cc forms a CA-GT-skew-associated local distribution pattern across the human genome

To identify features associated with TOP1cc positioning, we performed chromatin hidden Markov model (ChromHMM) analysis integrated TOP1cc, total TOP1 occupancy, and chromatin features. A 15-state genomic classification separated regions enriched for TOP1cc or TOP1 from another dominated by TOP1 binding, as well as from promoter, enhancer, transcribed, Polycomb-repressed, and heterochromatic states (Fig. 4A). Thus, local TOP1cc organization was not captured by total TOP1 occupancy or by any single conventional chromatin annotation.

**Fig. 4.**
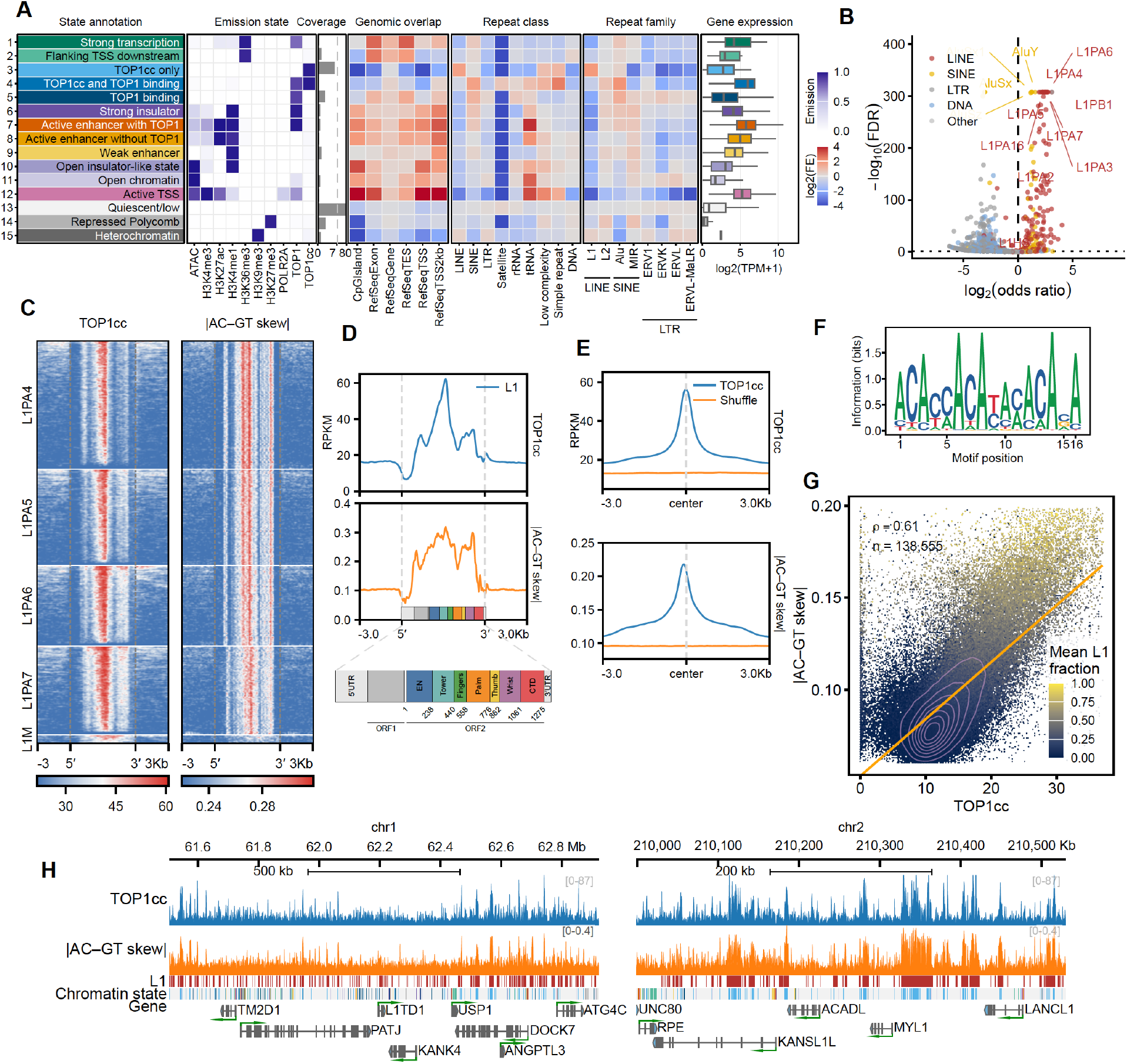
TOP1cc forms a sequence-context-dependent local distribution pattern across the human genome. **(A)** Fifteen-state genomic classification integrating chromatin features, TOP1 and TOP1cc coverage, genomic annotations, repeat classes and families, and gene expression. Colors in the emission-state matrix indicate relative feature enrichment; genomic-overlap values are shown as log2 fold enrichment. **(B)** Enrichment of repeat families in TOP1cc-associated regions; selected LINE-1 (L1) and other repeat families are labeled. **(C)** Heatmaps of TOP1cc signal and absolute CA–GT sequence skew across full-length L1PA4, L1PA5, L1PA6, L1PA7, and L1M elements with 3-kb flanking regions. **(D)** Average TOP1cc signal and absolute CA–GT skew across a full-length L1 element, aligned to the indicated L1 structural annotation. **(E)** Average TOP1cc signal and absolute CA–GT skew centered on TOP1cc sites and matched shuffled regions across ±3-kb flanking regions. **(F)** Sequence-information profile surrounding standard pDAIP-seq TOP1cc-associated positions. **(G)** Relationship between TOP1cc signal and absolute CA–GT skew across 138,555 genomic intervals (Spearman ρ = 0.61). Points are colored by mean L1 fraction. **(H)** Representative regions on chromosomes 1 and 2 showing TOP1cc signal, absolute CA–GT skew, L1 elements, chromatin states, and gene annotations.

We next asked what TOP1cc-associated regions share at the sequence level. Repeat enrichment analysis revealed strong enrichment of several LINE-1 families in TOP1cc-associated regions (Fig. 4, A and B). Alignment across full-length L1 elements showed reproducible internal TOP1cc enrichment in multiple L1PA subfamilies, accompanied by similarly positioned variation in absolute CA–GT skew (Fig. 4, C and D). TOP1cc-centered profiles likewise showed enrichment of absolute CA–GT skew relative to matched shuffled regions (Fig. 4E), and sequence-information analysis of standard pDAIP-seq identified a defined local sequence preference around TOP1cc-associated positions (Fig. 4F). Across 138,555 genomic intervals, TOP1cc signal correlated with absolute CA–GT skew, with the strongest values preferentially occurring in regions with greater L1 content (Fig. 4G). Representative genomic regions further showed correspondence among TOP1cc signal and absolute CA–GT skew (Fig. 4H).

Together, these analyses identify a strong association between TOP1cc positioning and local DNA sequence features, particularly the CA–GT skew. However, sequence composition can covary with repeat identity, chromatin state, transcription, GC content, and other genomic properties across the genome, so these cellular associations do not establish that primary DNA sequence itself contributes directly to TOP1cc positioning. To isolate this contribution from the surrounding cellular environment, we asked whether features of the TOP1cc landscape could be reconstructed on purified DNA templates.

### Local DNA sequence context is a major determinant of TOP1cc positioning

We developed an in vitro pDAIP-seq workflow (vipDAIP-seq) in which purified TOP1 was incubated with PCR-amplified genomic fragments and the resulting covalent complexes were processed using the same core enrichment strategy as cellular pDAIP-seq (Fig. 5A). PCR amplification preserves primary DNA sequence while removing chromatin, transcription, R-loops, DNA topology, and most other features of the cellular environment, providing a reduced system in which intrinsic sequence-dependent cleavage can be examined. We found that in vitro TOP1cc signal reproduced distributions and peaks of the cellular TOP1cc profiles across three genomic regions randomly picked from the human genome (Fig. 5B). Across 1-kb bins, cellular and in vitro signals were significantly correlated (Fig. 5C). Thus, part of the cellular TOP1cc pattern can be reconstructed from PCR-derived DNA in the absence of its chromatin and cellular context.

**Fig. 5.**
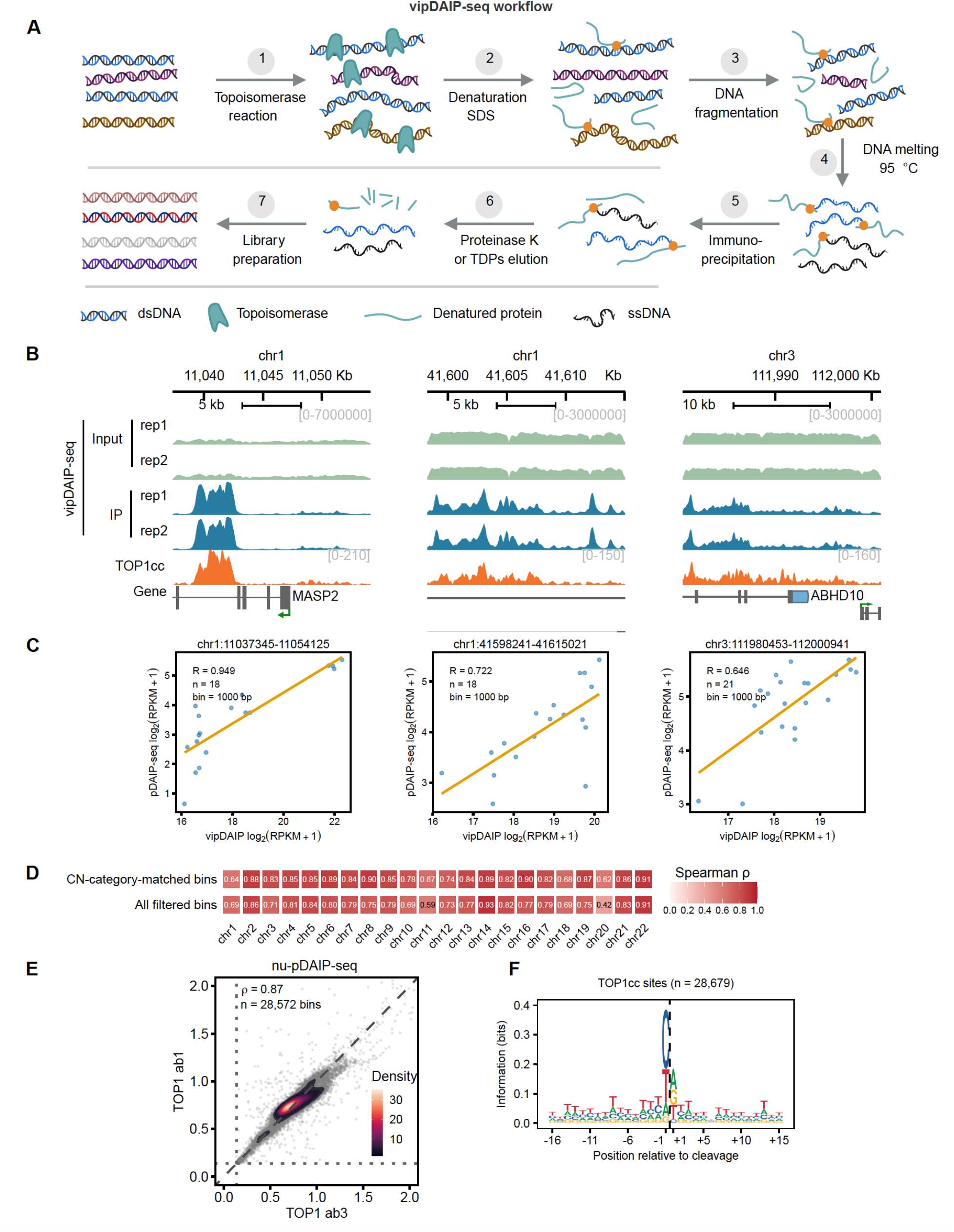
DNA sequence is a primary determinant of TOP1cc positioning. **(A)** Schematic of the in vitro pDAIP-seq (vipDAIP-seq) workflow. **(B)** Three representative genomic regions comparing vipDAIP-seq input and immunoprecipitated (IP) signal from two replicates with cellular TOP1cc signal and gene annotations. **(C)** Pearson correlations between cellular pDAIP-seq and vipDAIP-seq signal in 1-kb bins across the three regions shown in (B). Correlation coefficients are R = 0.949, 0.722, and 0.646 for 18, 18, and 21 bins, respectively. **(D)** Chromosome-level Spearman correlations between HCT116 and MCF-7 TOP1cc signal calculated using all filtered bins or bins matched by copy-number (CN) category. **(E)** Genome-wide comparison of nucleotide-resolution pDAIP-seq (nu-pDAIP-seq) signal generated with two TOP1 antibodies across 28,572 bins. **(F)** Sequence-information profile surrounding 28,679 nucleotide-resolved TOP1 cleavage sites; the dashed line marks the cleaved nucleotide.

As an orthogonal comparison, we compared TOP1cc profiles between HCT116 and MCF-7 cells after accounting for copy-number differences. Chromosome-level correlations remained significant when analysis was restricted to bins assigned to the same copy-number category in both cell lines (Fig. 5D), further supporting the contribution of DNA sequence context to TOP1cc positioning.

We further developed nucleotide-resolution pDAIP-seq (nu-pDAIP-seq), which maps the TOP1 cleavage site at nucleotide resolution. With reproducible genomic signals from independent TOP1 antibodies (Fig. 5E), nu-pDAIP-seq revealed the sequence context preference of the cleaved nucleotide (Fig. 5F), consistent with the CA–GT skew observed for TOP1cc genomic distribution.

Together with the cellular sequence associations, vipDAIP-seq and nu-pDAIP-seq support that primary DNA sequence context contains intrinsic information that contributes to genomic TOP1cc positioning. The incomplete reconstruction of the cellular landscape indicates that sequence is not sufficient by itself and that additional cellular features may also shape TOP1cc distribution. These results support a model in which topological demand regulates how much TOP1cc accumulates globally, whereas local sequence context contributes to where those complexes preferentially form.

### Genome-wide mapping of covalent intermediates involved in DNA repair reveals endogenous TOP1cc as a potential source of spontaneous DNA breaks

TOP1cc normally undergoes rapid religation, but occasionally formed abortive catalytic intermediates can generate trapped TOP1cc, leading to protein-linked DNA ends that require processing by DNA damage and repair systems in human cells. We therefore asked whether TOP1cc-rich genomic regions also coincide with DNA-end and repair-associated covalent chemistry. To test this possibility, we extended pDAIP-seq to TDP1, which forms a transient phosphohistidyl-DNA intermediate during processing of phosphotyrosyl DNA ends (*27*), DNA single-strand-break sensor PARP1, which can form covalent intermediates at 3′-polyunsaturated aldehyde (3′-PUA) termini (*28*, *29*), and DNA double-strand-break sensor component KU70, which can form a Schiff-base intermediate at abasic or deoxyribose-phosphate sites (*30*, *31*) (Fig. 6A). Interestingly, TOP1cc, TDP1, PARP1, and KU70 pDAIP-seq datasets were positively correlated across the genome, with varying degree of correlation coefficients (Fig. 6B). Similar patterns were also observed around genes between TOP1cc and these factors involved in DNA repair (Fig. 6C). We further included genome-wide signals of AP-seq and GLOE-seq, mapping abasic sites and 3′-hydroxyl DNA ends, respectively. We found that TDP1, PARP1, KU70, as well as AP-seq and GLOE-seq signals, were all enriched around endogenous TOP1cc sites relative to shuffled controls (Fig. 6D). Peak-centered heatmaps and a representative genomic region further confirmed the general correlations between them across the genome (Fig. 6, E and F).

**Fig. 6.**
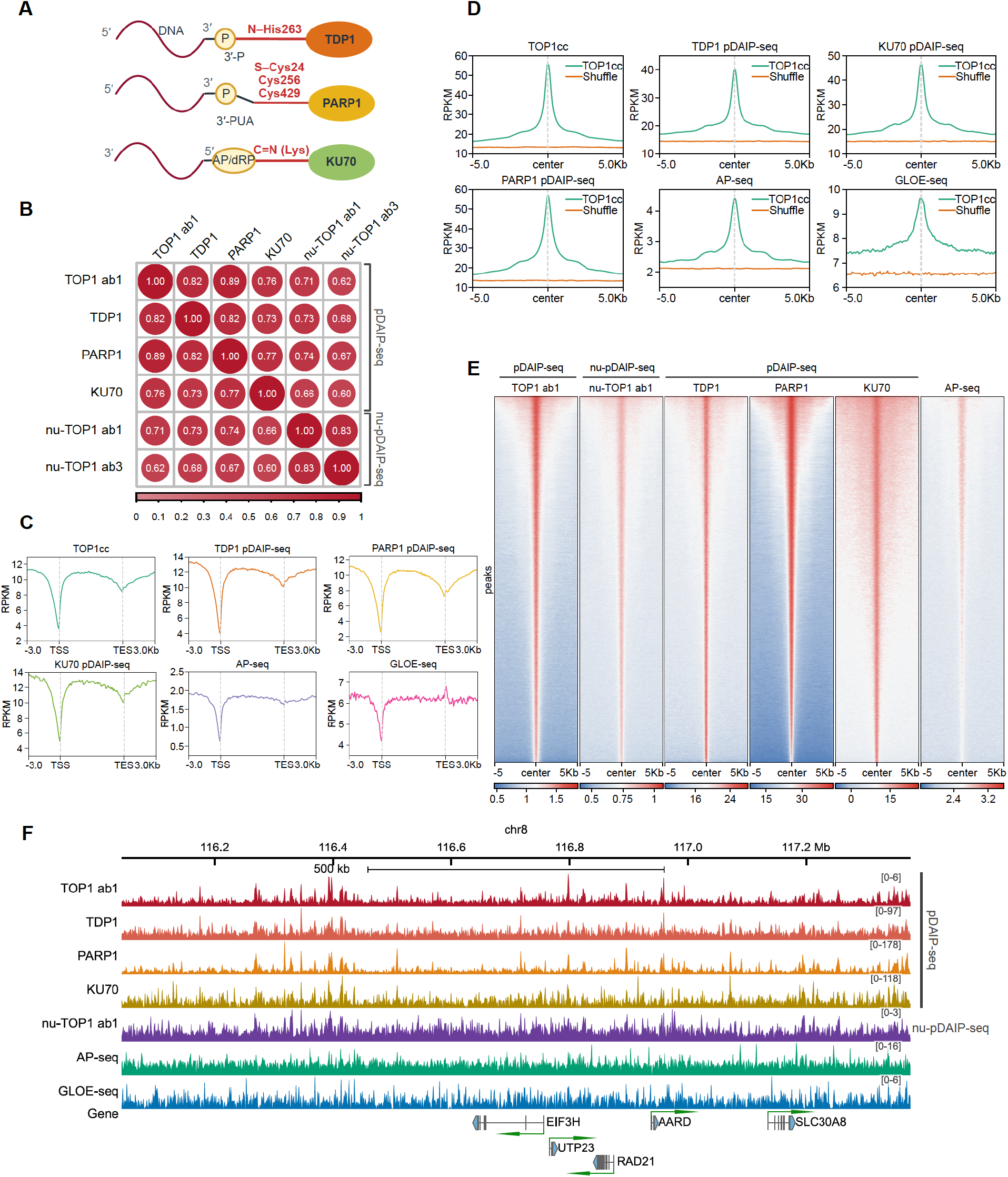
Endogenous TOP1cc is a potential major source of spontaneous DNA damage. **(A)** Schematics of the TDP1 phosphohistidyl intermediate at His263, cysteine-linked PARP1 intermediates at 3′-polyunsaturated aldehyde (3′-PUA) termini, and the lysine-mediated KU70 Schiff-base intermediate at 5′ deoxyribose-phosphate/abasic (5′-dRP/AP) sites. **(B)** Pairwise Spearman correlations among TOP1cc, TDP1, PARP1, and KU70 pDAIP-seq and nu-pDAIP-seq datasets generated with two TOP1 antibodies. **(C)** Gene-centered profiles of TOP1cc, TDP1, PARP1, and KU70 pDAIP-seq, AP-seq, and GLOE-seq signal. **(D)** Average signal from the indicated datasets centered on TOP1cc sites or matched shuffled regions across 5-kb flanking regions. **(E)** Heatmaps of TOP1cc, TDP1, PARP1, and KU70 pDAIP-seq, TOP1 nu-pDAIP-seq, and AP-seq signal centered on TOP1cc peaks across 5-kb flanking regions. **(F)** Representative region on chromosome 8 showing pDAIP-seq, nu-pDAIP-seq, AP-seq, GLOE-seq, and gene tracks. TDP1, tyrosyl-DNA phosphodiesterase 1; AP-seq, abasic-site sequencing; GLOE-seq, genome-wide ligation of 3′-hydroxyl ends followed by sequencing.

These relationships establish a spatial association between TOP1cc, DNA-repair-associated factors, and exposed DNA ends, indicating topoisomerase catalysis, particularly endogenous TOP1cc, as a potential major source of spontaneous DNA breaks in human cells.

### pDAIP-seq extends across topoisomerase classes and SPO11-DNA intermediates in the primary tissue

We next asked whether the pDAIP-seq principle was specific to TOP1 or could capture cleavage complexes formed by other topoisomerase classes. We profiled and compared the genome-wide patterns of TOP1cc, TOP2Acc, TOP2Bcc, TOP3Acc, and TOP3Bcc in human cells (Fig. 7, A, B, and D). Intersection analysis identified both shared and enzyme-specific genomic regions (Fig. 7C). Pairwise correlations further resolved two broad relationships: TOP1cc was most similar to TOP3Bcc and TOP3Acc, and TOP2Acc was most similar to TOP2Bcc, while correlations between these groups were lower (Fig. 7E). These results demonstrate that protein-specific immunoprecipitation of pDAIP-seq retains enzyme identity even among related covalent cleavage chemistries.

**Fig. 7.**
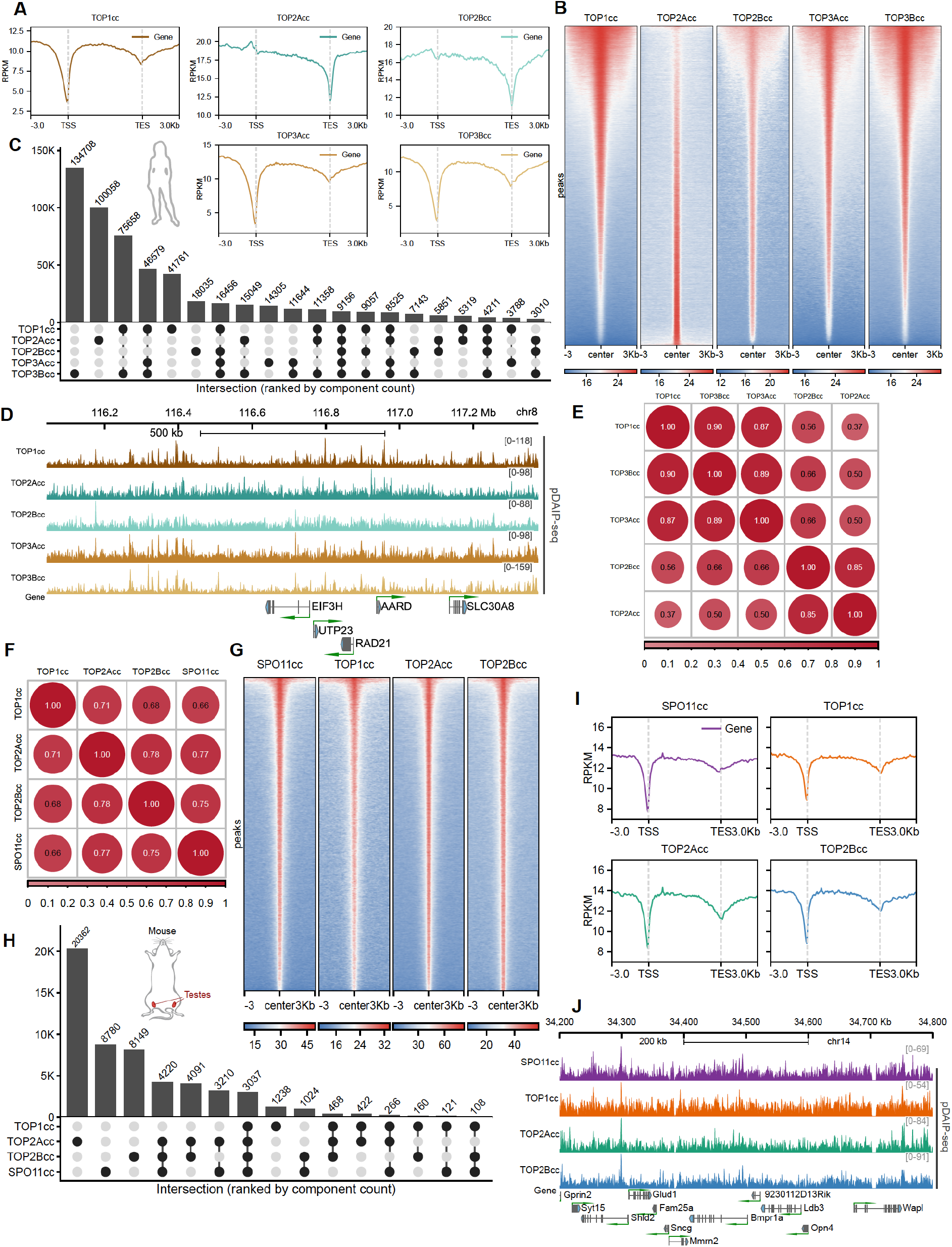
pDAIP-seq extends across topoisomerase classes and in primary tissues. **(A)** Gene-centered profiles of human TOP1cc, TOP2Acc, TOP2Bcc, TOP3Acc, and TOP3Bcc signals across transcription units and 3-kb flanking regions. **(B)** Heatmaps of the five human topoisomerase cleavage-complex signals centered on their union peaks with 3-kb flanking regions. **(C)** Intersection analysis of human TOP1cc, TOP2Acc, TOP2Bcc, TOP3Acc, and TOP3Bcc regions, ranked by intersection size. **(D)** Representative region on chromosome 8 showing the five human topoisomerase cleavage-complex pDAIP-seq tracks and gene annotations. **(E)** Pairwise Spearman correlations among the five human topoisomerase cleavage-complex datasets. **(F)** Pairwise Spearman correlations among mouse-testis TOP1cc, TOP2Acc, TOP2Bcc, and SPO11cc. **(G)** Heatmaps of mouse-testis SPO11cc, TOP1cc, TOP2Acc, and TOP2Bcc signals centered on their union peaks with 3-kb flanking regions. **(H)** Intersection analysis of mouse-testis TOP1cc, TOP2Acc, TOP2Bcc, and SPO11cc regions; the inset indicates the tissue source. **(I)** Gene-centered profiles of mouse-testis SPO11cc, TOP1cc, TOP2Acc, and TOP2Bcc signals across transcription units and 3-kb flanking regions. **(J)** Representative region on mouse chromosome 14 showing SPO11cc, TOP1cc, TOP2Acc, TOP2Bcc, and gene tracks. TSS, transcription start site; TES, transcription end site; RPKM, reads per kilobase per million mapped reads.

We then asked whether pDAIP-seq could recover a physiological, developmentally programmed covalent DNA intermediate directly from primary tissues. We picked mouse testis, in which SPO11 creates programmed meiotic DNA double-strand breaks through a covalent 5′-phosphotyrosyl intermediate that initiate meiotic recombination and was recently shown to also catalyze reversible single-strand cleavage and religation in vitro (*32*, *33*). These findings indicate that SPO11 can populate transient covalent DNA intermediates, raising the question of how such intermediates are distributed across the genome without exogenous poison treatment. Moreover, tissue-expression and protein-abundance datasets also suggested the presence of the topoisomerases and potentially TOPcc in testis (fig. S5). Using pDAIP-seq, we recovered SPO11-associated signal together with TOP1cc, TOP2Acc, and TOP2Bcc genomic profiles in mouse testis. They shared broad genomic patterns but remained distinguishable (Fig. 7, F and H), particularly underscore the similar genome-wide distribution between SPO11-associated intermediates and topoisomerase cleavage complexes in mouse testis, from peak-centered analysis, gene-region analysis, and an example genomic region (Fig. 7, G, I, and J).

## Discussion

Transient covalent protein–DNA intermediates are intrinsic to topoisomerase catalysis and also occur during DNA repair and programmed DNA cleavage, but their short lifetimes complicate their capture and genomic analysis. pDAIP-seq combines the recovery of covalently protein-linked DNA with protein-specific immunoprecipitation, allowing these complexes to be mapped without prior poison stabilization while retaining the identity of the linked protein.

pDAIP-seq addresses two features of topoisomerase biochemistry that make endogenous TOPcc difficult to capture. First, the catalytic intermediate is chemically reversible: topoisomerases retain the capacity to cleave and religate DNA under permissive conditions, so preservation of TOPcc requires minimizing catalytic reversal during isolation (*18*, *19*, *32*, *34–36*). Second, topoisomerases can participate in RNA-associated interactions—TOP1 modulates DNA topology through direct RNA binding (*37*), RNA regulates TOP2A and TOP2B catalytic activity (*38*, *39*), and TOP3B participates in ribonucleoprotein complexes (*40*)—while proteomic analyses further reveal diverse interaction networks for human topoisomerases (*41*). These properties create potential routes for noncovalent recovery that can obscure the covalent population. The pDAIP-seq workflow was accordingly designed with denaturing extraction and RNase treatment to reduce noncovalent protein and RNA carry-over, and with DNA melting immediately before immunoprecipitation to disrupt the duplex geometry required for TOP1 religation. Selective enrichment of covalent TOP1cc was further supported by IgG and proteinase K controls and by the catalytically inactive GFP-TOP1^Y723F^ mutant.

These design features distinguish pDAIP-seq from existing approaches in complementary ways rather than through a direct performance comparison, which we did not attempt. Non-crosslinking ChIP-like enrichment may remain susceptible to residual noncovalent interactions and catalytic reversal during immunoprecipitation, potentially blurring the distinction between total topoisomerase occupancy and the covalent intermediate. TOP1 CAD-seq uses MG132 and CPT to stabilize TOP1cc (*12*), whereas pDAIP-seq is designed to recover the untreated population. CC-seq achieves nucleotide-resolution mapping of endogenous TOP2cc and SPO11 complexes but relies on chemistry-based enrichment that does not itself encode the identity of the linked protein (*15*). pDAIP-seq therefore contributes the combination of poison-free recovery and antibody-defined protein specificity, enabling the same general strategy to be applied across different topoisomerases, repair-associated adducts, and primary tissue.

The separation between total TOP1 occupancy and TOP1cc suggests that recruitment and catalytic engagement are distinct regulatory steps. TOP1 has been shown to enhance assembly of the TFIID–TFIIA complex during transcriptional activation, and this coactivator activity can be retained by catalytically inactive TOP1 (*42*). Thus, abundant promoter-associated TOP1 need not correspond to proportionate accumulation of the covalent cleavage intermediate. Catalytic engagement, by contrast, should depend on the topological state encountered by the enzyme. The reciprocal responses of TOP1cc to transcriptional suppression and TOP2A depletion are consistent with genome-wide topological demand influencing the fraction of TOP1 that enters the covalent catalytic state, rather than TOP1cc abundance being determined solely by local TOP1 occupancy or transcription level.

This framework further suggests that the determinants of how much TOPcc forms need not be identical to those determining where cleavage occurs across the genome in human cells. Using SHAN-seq *in vitro*, Morgan et al. found that DNA supercoiling and supercoil chirality altered the overall frequency of bacterial gyrase and topoisomerase IV cleavage complexes while having comparatively little effect on cleavage-site specificity, whereas ciprofloxacin substantially changed both the location and relative frequency of cleavage complexes (*43*). Although bacterial type II topoisomerases differ substantially from human TOP1, these findings provide an independent *in vitro* biochemical precedent for topology primarily tuning cleavage-complex abundance while local substrate properties contribute more strongly to cleavage positioning. They also reinforce the broader caution that topoisomerase poisons can reshape cleavage-complex distributions rather than simply amplify an otherwise unchanged population.

Local DNA sequence composition provides a second, complementary layer. Sequence preference at the cleaved nucleotide, its relationship with regional absolute CA–GT skew, and partial reconstruction of the cellular pattern on purified DNA together indicate that primary DNA sequence carries intrinsic positional information but does not fully specify the cellular TOP1cc landscape. Beyond mapping, vipDAIP-seq provides a controlled biochemical framework in which sequence variants, DNA lesions, altered DNA-end chemistry, DNA topology, or modified TOP1 proteins can be introduced individually to determine how each affects covalent-intermediate formation. Compared with cleavage-end–based approaches such as SHAN-seq, vipDAIP-seq directly enriches the covalent protein–DNA adduct by antibody before library preparation rather than identifying complexes through processed DNA ends, thereby depleting unreacted DNA and potentially increasing effective sensitivity for low-abundance complexes on large or complex substrates. Because the same protein-defined capture principle is used in cellular pDAIP-seq, purified and cellular systems can be compared directly, and the strategy can be extended across topoisomerase classes and to other covalent protein–DNA intermediates.

The distinction between covalent and end-defined lesions has broader implications for how endogenous DNA damage is detected. Oxidative lesions, abasic sites, and other spontaneous damage are well-established sources of endogenous strand breaks, but an abortive TOP1 catalytic cycle produces a distinct lesion in which the DNA end remains blocked by covalently attached TOP1 and may therefore be poorly accessible to assays that require a free or chemically compatible DNA terminus until TOP1 is proteolyzed or its phosphotyrosyl linkage is processed(*5*, *44*, *45*). The genomic association of TOP1cc with TDP1, PARP1, and KU70 covalent-adduct maps, together with AP-seq and GLOE-seq signal around TOP1cc-rich regions, is consistent with the possibility that a fraction of physiological TOP1 cleavage intermediates fails to religate efficiently and enters DNA-end processing pathways. The present data do not fully establish that these repair-associated intermediates, including TDP1, PARP1, and KU70, originate directly from TOP1cc, since the observed relationship is correlation rather than causal. Nevertheless, the results raise the possibility that DNA lesions generated during abortive TOP1 catalysis under normal physiology may be one of the major endogenous sources of human genome instability(*46*). Testing this model will require perturbing TOP1cc formation or processing and determining whether repair-associated covalent intermediates and DNA-end signals change accordingly.

Extending pDAIP-seq beyond TOP1 broadens this framework. Protein-defined mapping of TOP1cc, TOP2Acc, TOP2Bcc, TOP3Acc, and TOP3Bcc provides a basis for investigating how catalytic mechanism, protein partners, chromatin context, RNA-associated regulation, and DNA topology differentiate the activities of individual topoisomerases. SPO11 offers a particularly informative application. SPO11 initiates meiotic recombination through a covalent protein–DNA intermediate (*33*), and recent biochemical work indicates that it can also catalyze reversible single-strand cleavage and religation (*32*), raising the possibility that meiotic cells contain a broader spectrum of SPO11-associated covalent states than the processed double-strand-break products measured by conventional SPO11-oligonucleotide approaches(*47*). pDAIP-seq provides a complementary means of recovering SPO11-associated covalent DNA directly from primary tissues. Because the present measurements were obtained from bulk testis and do not distinguish among all intact and processed SPO11-associated species, developmental and genetic perturbations will be required to define which meiotic intermediates contribute to the observed signal.

Several considerations bound the interpretation of pDAIP-seq. The assay measures covalent complexes recoverable at the time of isolation and therefore reflects the combined effects of formation, reversal, and processing rather than catalytic flux or the intrinsic rates of supercoiling relaxation. DNA melting increased TOP1cc recovery in a manner consistent with reduced post-isolation reversal but does not establish the extent to which reversal is prevented. Standard pDAIP-seq maps enriched protein-linked fragments rather than the exact covalent bond, which requires the end-resolved nu-pDAIP-seq implementation. Recovery also depends on antibody performance, limiting absolute quantitative comparisons across different proteins. Because pDAIP-seq measurements represent population averages, they cannot distinguish changes in the fraction of catalytically engaged TOP1 from changes in the lifetime of individual TOP1cc.

Taken together, our findings establish pDAIP-seq as a broadly applicable framework for studying covalent protein–DNA intermediates under normal physiological conditions. For TOP1 specifically, the data support a model in which poison stabilization remodels the endogenous landscape, genome-wide maintenance of DNA topological homeostasis regulates the abundance of endogenous TOP1cc, and local DNA sequence contributes to their genomic positioning. The genome-wide association of TOP1cc with repair-associated covalent complexes and endogenous DNA breaks further suggests endogenous TOP1cc at the interface between DNA topological regulation, DNA damage/repair, and human genome instability.

## Acknowledgments

We thank Qin Li (Laboratory of Human Carcinogenesis, NCI, NIH) for helping collect mouse testis tissues, Kazuhiro Maeshima (National Institute of Genetics, Japan) for DLD1 Tet-OsTIR1/mAID-mClover-POLR2A cell line, Laura Baranello (Karolinska Institute) and Christian Friberg Nielsen (Copenhagen University) for HCT116 CMV-OsTIR1/TOP2A-mAID H2B-eGFP cell line, Federico Machinandiarena (Laboratory of Molecular Biology, NCI, NIH) for MCF-7 cell line. We thank Madeline Wong and Steve Shema from the CCR Genomics Core for sequencing. This study used the Biowulf Linux cluster at the National Institutes of Health. Figures were created using Biorender.com.

## Funding

This work was supported by the Intramural Research Program of the National Institutes of Health, National Cancer Institute (1ZIABC011884). The contributions of the NIH authors were made as part of their official duties as NIH federal employees, are in compliance with agency policy requirements, and are considered Works of the United States Government. However, the findings and conclusions presented in this paper are those of the authors and do not necessarily reflect the views of the NIH or the U.S. Department of Health and Human Services.

## Author contributions

S.L. and C.C. conceived the study. S.L. developed the methodology, performed the experimental work and data analysis, and prepared the visualizations. C.C. administered the project and supervised the study. S.L. and C.C. wrote the manuscript.

## Competing interests

Authors declare that they have no competing interests.

## Data, code, and materials availability

All data, code, and materials used in the analysis reported in this paper is available from the corresponding author upon request. This paper does not report original code or generate new unique reagents.

## Supplementary Materials

Materials and Methods

Figs. S1 to S5

## Notes

### Competing Interest Statement

The authors have declared no competing interest.

