## supplemental tables and figures for "Genome-wide mapping of endogenous topoisomerase cleavage complexes reveals TOP1 at the interface of DNA topological regulation and genome instability"

### Supplementary Materials

#### Materials and Methods

##### Cell culture and chemical treatments

HCT116 and DLD1 cells were maintained in McCoy's 5A Modified Medium, and MCF-7 cells were maintained in DMEM. Cells were maintained at 37 °C in a humidified incubator with 5% CO2. All cultures were supplemented with 10% fetal bovine serum and penicillin-streptomycin. Where indicated, cells were treated with 20 µM camptothecin (CPT) for 5 min, 2 µM flavopiridol for 1 h, or 1 µM triptolide for 1 h.

##### Degron cell lines and acute protein depletion

POLR2A depletion experiments used DLD1 Tet-OsTIR1/mAID-mClover-POLR2A cells described previously(*1*). Cells were treated with 2 µg/ml doxycycline for 48 h to induce OsTIR1 and then with 500 µM 1-naphthaleneacetic acid (NAA) for 4 h to degrade mAID-mClover-POLR2A. DMSO control cells did not add doxycycline. Depletion was confirmed by immunoblotting before genomic analysis.

TOP2A depletion experiments used HCT116 CMV-OsTIR1/TOP2A-mAID H2B-eGFP cells described previously(*2*). Control cells were treated with 100 µM auxinole for 24 h, whereas TOP2A-depleted cells were treated with 500 µM NAA for 4 h. Depletion was confirmed by immunoblotting before genomic analysis.

##### Immunoblotting

Cells were lysed in 2× RIPA buffer and incubated on ice for 20 min. Lysates were diluted to 1× RIPA and treated with Benzonase nuclease for 20 min on ice. After clarification, protein concentrations were determined using DC™ Protein Assay Kit II, and 10–20 µg of total protein was resolved by SDS-PAGE and analyzed by immunoblotting. Primary antibodies were diluted 1:3000 and incubated for 1 h at room temperature; HRP-conjugated secondary antibodies were diluted 1:5000 and incubated for 1 h at room temperature.

##### Chromatin immunoprecipitation sequencing

ChIP-seq were performed as previously described with small modifications(*3*). ChIP-seq Cells were crosslinked with 1% formaldehyde for 10 min at room temperature and quenched with glycine at a final concentration of 0.125 M for 5 min. Cells were collected at 800 × g for 5 min at 4 °C, washed twice with ice-cold PBS, divided into aliquots of 7–10 million cells, and stored at −80 °C. Crosslinked cells were lysed for 5 min on ice in 0.5 ml NP-40 lysis buffer (10 mM Tris-HCl, pH 7.5, 150 mM NaCl, and 0.05% NP-40). Lysates were layered over 1.25 ml of 24% sucrose in NP-40 lysis buffer and centrifuged at 13,000 × g for 10 min at 4 °C. Nuclei were washed once with PBS containing 1 mM EDTA.

Nuclei were resuspended in 0.5 ml glycerol buffer (20 mM Tris-HCl, pH 8.0, 75 mM NaCl, 0.5 mM EDTA, 0.85 mM DTT, and 50% glycerol) and mixed with 0.5 ml nuclear lysis buffer (10 mM HEPES, pH 7.6, 1 mM DTT, 7.5 mM $MgCl_2$, 0.2 mM EDTA, 0.3 M NaCl, 1 M urea, 1% NP-40, and 1% Empigen BB) for 2 min on ice. After centrifugation at 13,000 × g for 2 min, the chromatin pellet was washed twice with PBS containing 1 mM EDTA and resuspended in 1 ml sonication buffer (20 mM Tris-HCl, pH 8.0, 150 mM NaCl, 0.1% SDS, 1% Triton X-100, and 1% Empigen BB). Chromatin was digested with 400 U micrococcal nuclease in 5 mM $CaCl_2$ for 15 min at 37 °C with shaking at 700 rpm. The reaction was stopped on ice by adding 20 µl of 0.5

M EDTA and 40 µl of 0.5 M EGTA. Chromatin was then sheared for 6 min using Covaris M220 using milliTUBE 1 ml AFA Fiber at a peak incident power of 75 W, 15% duty factor, and 200 cycles per burst.

For TOP1 ChIP-seq, 2 µg of TOP1 antibody was added to cleared chromatin and incubated overnight at 4 °C with rotation. Prewashed Pierce Protein A/G Magnetic Beads were added and incubated for 4 h. Beads were washed twice with sonication buffer, twice with high-salt buffer (20 mM Tris-HCl, pH 8.0, 500 mM NaCl, 2 mM EDTA, 0.1% SDS, and 1% Triton X-100), twice with LiCl wash buffer (10 mM Tris-HCl, pH 8.0, 250 mM LiCl, 1 mM EDTA, and 1% NP-40), and twice with TE. Beads were transferred to a new tube before the final wash. DNA eluted in 30 µl of TE containing proteinase K at a final concentration of 100 µg/ml overnight at 65 °C, followed by 10 min at 95 °C to denature proteinase K. DNA was used directly for library preparation or, when needed, purified with 2.5 volumes of SPRIselect beads before library preparation with the xGen ssDNA and Low-Input DNA Prep Kit.

#### Protein–DNA adduct immunoprecipitation sequencing

##### *pDAIP-seq*

Seven to ten million cells were either washed once with ice-cold PBS and immediately placed in an ice-water bath or add DNAzol directly after dumping the medium. Genomic DNA was isolated with DNAzol Reagent. After ethanol washing, the pellet was briefly air-dried, dissolved in 100 µl of TE (10 mM Tris-HCl, pH 8.0, and 1 mM EDTA) containing 1% SDS, and incubated for 10 min at room temperature. The sample was mixed with 100 µl of TE containing 100 ng/µl RNase A for 10–20 min at room temperature. DTT was then added to a final concentration of 10 mM, and the sample was incubated for an additional 10 min at 65 °C. The mixture was brought to 600 µl with nuclease-free water. Ammonium acetate was added to a final concentration of 0.75 M, followed by an equal volume of 2-propanol. DNA was collected at 4,000 × g for 5 min, washed with 70% ethanol, and dissolved in TE containing 0.5% SDS. DNA concentration was measured with the Qubit dsDNA Broad Range Assay kit.

Twenty to sixty micrograms of genomic DNA was sheared with a Covaris M220 using 130-µl AFA microtubes. DNA was fragmented to approximately 500 bp for most pDAIP-seq experiments and approximately 800 bp for TOP2A and TOP2B pDAIP-seq. For the melted preparation, DNA was heated at 95 °C for 5 min and immediately cooled on ice-water bath before immunoprecipitation; this step was omitted for the unmelted DNA comparison. The fragmented DNA was adjusted to 200 µl with 1× PBS containing 1% Triton X-100. Separately, a 300-µl antibody mixture containing 1–2 µg of target-specific antibody and, where applicable, ChIP Spike-in Chromatin and the corresponding ChIP Spike-in Antibody was prepared. For samples within the same direct experimental comparison, this component was prepared as a master mix and aliquoted equally to minimize pipetting variation in antibody and spike-in amounts. The 200-µl DNA fraction and 300-µl antibody mixture were combined to give a final 500-µl immunoprecipitation reaction. For TOP1 pDAIP-seq, the antibody mixture contained 8 ng of spike-in chromatin and 1 µg of spike-in antibody per immunoprecipitation. Spike-in amounts were empirically optimized for each target antibody; batch-specific amounts were used but held constant among samples within each direct experimental comparison.

The next day, 10–20 µl of Pierce Protein A/G Magnetic Beads was added and incubated for 3 h at 4 °C. Beads were washed twice for 7 min with IP buffer; twice for 7 min with 50 mM Tris-HCl, pH 8.0, 500 mM NaCl, 1 mM EDTA, 1% Triton X-100, and 0.1% SDS; twice for 7 min with 10 mM Tris-HCl, pH 8.0, 250 mM LiCl, 1 mM EDTA, 1% NP-40, and 1% sodium deoxycholate; and twice for 5 min with TE. Beads were transferred to a new tube before the final wash.

DNA was eluted in 30 µl of TE containing proteinase K at a final concentration of 100 µg/ml, prepared from a 20 mg/ml stock, and incubated for 2 h at 55 °C, followed by 10 min at 95 °C. Following proteinase K digestion, samples were treated for 1 h with 0.5 µl of human TDP1 for TOP1, PARP1, and KU70 pDAIP-seq or with 0.5 µl of human TDP2 for TOP2A, TOP2B, TOP3A, TOP3B, and SPO11 pDAIP-seq in the corresponding 1× reaction buffer in a final volume of 55 µl. No TDP1 or TDP2 treatment was applied to TDP1 pDAIP-seq samples. For standard pDAIP-seq, recovered DNA was sheared to approximately 250 bp in a 55-µl AFA microtube and purified with 2.5 volumes of SPRIselect beads. We have tested that these library kits were compatible to unpurified products without losing the efficiency. Libraries were prepared using either the VAHTS ssDNA Library Prep Kit for Illumina or the xGen ssDNA and Low-Input DNA Prep Kit, as used for the corresponding experiment.

Unless otherwise indicated, pDAIP-seq of TDP1, PARP1, KU70, TOP2A, TOP2B, TOP3A, TOP3B, and SPO11 followed the standard pDAIP-seq procedure described above with target-specific antibodies listed in table S1.

**pDAIP-seq validation and control experiments**

Doxycycline-inducible N-terminal GFP-TOP1 and GFP-TOP1$^{Y723F}$ HCT116 cell lines were generated in-house by integration of the corresponding constructs at the AAVS1 locus, with endogenous TOP1 left intact. The TOP1$^{Y723F}$ catalytic-site mutation was introduced by PCR-based mutagenesis. Expression was induced with 2 µg/ml doxycycline for 48 h.

For pDAIP-seq specificity controls, rabbit IgG was used at 2 µg per immunoprecipitation. For proteinase K pretreatment, genomic DNA was resuspended in 1× TE containing 0.5% SDS and proteinase K at a final concentration of 100 µg/ml and incubated overnight at 37 °C before immunoprecipitation. DNA-melting controls were performed by omitting the 95 °C melting step before immunoprecipitation. Method-development experiments additionally compared 5, 10, and 50 µg genomic DNA input and library amplification using 8 or 13 PCR cycles.

**vipDAIP-seq**

PCR-amplified genomic fragments were used as protein-free substrates for the in vitro pDAIP-seq assay (vipDAIP-seq). Genomic DNA extracted using the pDAIP-seq procedure was used as PCR template, with 200 ng genomic DNA per amplification. Target regions were amplified using LongAmp Taq 2X Master Mix or KOD One PCR Master Mix with locus-specific primer pairs listed in table S3. PCR products were purified with SPRIselect beads.

For the TOP1 cleavage reaction, 200 ng of each of three purified PCR amplicons was combined (600 ng total DNA) and incubated with 300 ng purified human TOP1. A 20-µl reaction contained the pooled PCR-amplified DNA, 2 µl of 10× TOP1 cleavage buffer (100 mM Tris-HCl, pH 7.5, 500 mM KCl, 50 mM $MgCl_2$, 1 mM EDTA, and 150 µg/ml BSA), purified human TOP1, and nuclease-free water. Reactions were incubated for 15 min at 25 °C and stopped with 0.5 µl of 20% SDS. DNA purification was omitted after the reaction, and samples proceeded directly to sonication and the remaining pDAIP-seq workflow. The three main vipDAIP-seq regions shown in Fig. 5 correspond to chr1:11,037,345–11,054,125 (MASP2), chr1:41,598,241–41,615,021 (HIVEP3), and chr3:111,980,453–112,000,941 (ABHD10).

**nu-pDAIP-seq**

Nucleotide-resolution pDAIP-seq (nu-pDAIP-seq) followed the standard pDAIP-seq procedure with two modifications. DNA was sheared to approximately 300 bp before immunoprecipitation, and the second sonication step before library preparation was omitted. Following immunoprecipitation, TOP1-linked DNA ends were released with proteinase K and then human TDP1. Libraries were prepared with ssDNA Prep Kit so that the DNA end corresponding to the covalent TOP1–DNA junction was retained. The first aligned genomic nucleotide of read 2 was used as the exact TOP1 cleavage coordinate.

**DRIP-Rx**

Genomic DNA was isolated with DNAzol. Residual proteins were removed by resuspending the DNA in 500 µl of 1× TE containing 0.5% SDS and 1 µg proteinase K (2 µg/ml) and incubating at 37 °C overnight. Protein debris was removed by adding a final concentration of 0.75 M ammonium acetate followed by 500 µl of 2-propanol. The DNA pellet was washed twice with 70% ethanol, air-dried, and resuspended in 150 µl TE.

To reduce nonhybrid RNA species that can contribute to S9.6 binding while retaining RNA–DNA hybrids, samples were treated according to a previously described protocol(*4*). Each sample received 2 µl RNase T1, 2 µl ShortCut RNase III, 20 µl of 10× RNase treatment buffer (500 mM Tris-HCl, pH 7.5, 750 mM KCl, and 30 mM $MgCl_2$), and 26 µl nuclease-free water. Samples were incubated for 1 h at 37 °C, brought to 500 µl with TE, precipitated as described above, and resuspended in 200 µl TE. DNA was sonicated to approximately 500 bp in a 130-µl tube. For the specificity control, 40 µg sonicated DNA was treated overnight at 37 °C in a 100-µl reaction containing 10 µl of 10× RNase H buffer and 5 µl RNase H.

For each immunoprecipitation, 20 µg DNA was used, and 0.2 µg DNA was retained as input. The RNase H-treated control was also immunoprecipitated using 20 µg DNA. Each immunoprecipitation received 60 ng ChIP Spike-in Chromatin, 1 µg ChIP Spike-in Antibody, and 2 µg S9.6 antibody. Immunoprecipitation reactions were assembled and incubated using the same procedure as pDAIP-seq, except that DRIP-Rx used 1× PBS containing 0.1% Triton X-100 as the IP buffer. The following day, 20 µl Pierce Protein A/G Magnetic Beads was added and incubated for 3 h at 4 °C. Beads were washed three times with DRIP-Rx high-sodium wash buffer 10 mM Tris-HCl, pH 8.0, 300 mM NaCl, 1 mM EDTA, and 0.1% Triton X-100 to reduce nonspecific background while preserving RNA–DNA hybrid binding(*5*), followed by three washes with 1× PBS, 1 mM EDTA, and 0.1% Triton X-100. Elution and DNA recovery followed the pDAIP-seq procedure but without TDPs treatment. Recovered DNA was converted into sequencing libraries with the VAHTS ssDNA Library Prep Kit for Illumina.

**KAS-seq**

Cells were incubated in prewarmed complete culture medium containing 1 mM N3-kethoxal for 10 min at 37 °C. Genomic DNA was isolated with DNAzol and resuspended in TE containing 0.5% SDS and RNase A, followed by incubation for 20 min at 37 °C. Ammonium acetate was added to a final concentration of 0.75 M, followed by an equal volume of 2-propanol to precipitate the DNA.

For biotinylation, 2–3 µg genomic DNA was adjusted to 85 µl in 25 mM $K_3BO_3$, mixed with 10 µl of 10× PBS and 5 µl of 20 mM DBCO-PEG4-biotin, and incubated for 1.5 h at 37 °C with shaking at 500 rpm. Biotinylated DNA was purified, and 1 µg DNA was sonicated to approximately 500 bp.

Biotinylated DNA was captured with 5 µl Dynabeads MyOne Streptavidin C1 using 2× B&W buffer (10 mM Tris-HCl, pH 7.4, 1 mM EDTA, 2 M NaCl, and 0.1% Tween-20) for binding. Beads were washed five times with 1× B&W buffer (5 mM Tris-HCl, pH 7.4, 0.5 mM EDTA, 1 M NaCl, and 0.05% Tween-20). Enriched DNA was released by heating the beads in water at 95 °C for 10 min, which also removed the N3-kethoxal modification. Recovered DNA was converted into sequencing libraries with the VAHTS ssDNA Library Prep Kit for Illumina.

**Mouse testis pDAIP-seq**

Testes were collected from 8-week-old C57BL/6J mice, immediately frozen, and homogenized under liquid nitrogen. Two biological replicates were analyzed, with testes from three mice pooled per replicate. Genomic DNA was isolated and processed using the standard pDAIP-seq workflow described above, using 60 µg genomic DNA per immunoprecipitation. TOP1, TOP2A, TOP2B, and SPO11 covalent complexes were enriched with the corresponding target-specific antibodies. The animal study protocol (LHC-006-3) was reviewed and approved by the National Cancer Institute-Bethesda Animal Care and Use Committee (Bethesda, MD, USA).

**Published genomic datasets**

Published datasets analyzed in this study included HCT116 AP-seq (GSM4224428; GSE142284)(*6*), HCT116 G4P ChIP-seq (GSM7670068; GSE239694)(*7*), HCT116 WT control-siRNA GLOE-seq data (GSE134224/GSE134225)(*8*), DNA-supercoiling data from SRA experiments SRX21313606 and SRX21313607 (PRJNA1003820)(*9*), and RNA-seq data from GSE285698(*10*). HCT116 chromatin-feature datasets used for genomic-state analysis were obtained from ENCODE as peak BED files, including ATAC-seq (ENCFF107FCE), H3K4me1 (ENCFF839ERU), H3K4me3 (ENCFF698BEK), H3K9me3 (ENCFF787WUK), H3K27ac (ENCFF949DQN), H3K27me3 (ENCFF943RFI), H3K36me3 (ENCFF094STC), and POLR2A (ENCFF848IHI). Data were obtained from the corresponding public repositories and analyzed on the hg38 reference genome as appropriate. RNA-seq measurements were used to assign genes to the expression groups shown in the figures. Normal-tissue RNA-expression data were obtained from GTEx Analysis V11, and normal-tissue immunohistochemistry and mass-spectrometry protein-abundance data were obtained from the Human Protein Atlas.

**Sequencing-data processing and normalization**

For standard human pDAIP-seq, paired-end reads were adapter- and quality-trimmed with Trim Galore v0.6.10 using Phred+33, stringency 10, a minimum retained read length of 20 nt, and a maximum of 10 ambiguous bases. The first 10 nt from the 5′ end of read 2 were subsequently removed with Cutadapt v4.9 using -U 10 -m 20. Reads were aligned to hg38 or a combined hg38 and dm6 reference using Bowtie2 v2.5.4 with --very-sensitive-local, --no-mixed, --no-discordant, and --maxins 1000. PCR duplicates were removed using Picard MarkDuplicates v2.27.5 with REMOVE_DUPLICATES=true. Alignments with MAPQ ≥10 were retained using SAMtools v1.16.1.

Retained dm6 read counts were obtained with Spiker v1.0.3(*11*). Sample-specific spike-in scaling factors were calculated as 1,000,000 divided by the number of retained dm6 reads. Spike-in-normalized bigWig tracks were generated with deepTools v3.5.5 at 5-bp resolution using the corresponding scaling factor without additional library-size normalization. Spike-in-normalized signal was used for primary quantitative comparisons between experimental conditions. Spike-in normalization was interpreted within direct experimental comparisons in which the spike-in chromatin and antibody amounts were held constant. RPKM-normalized tracks were generated separately where RPKM is explicitly indicated in the figures. The effective human genome size used for coverage normalization was 2,747,877,702 bp.

**Peak calling**

Peaks were called from MAPQ-filtered, duplicate-removed BAM files using MACS3 v3.0.2 in paired-end mode (-f BAMPE) with --keep-dup all, --nomodel, --nolambda, and -q 0.1. Human peaks overlapping at least 25% of the ENCODE hg38 DAC exclusion list (ENCFF356LFX) were removed with BEDTools v2.31.1. For downstream analyses, the TDP1 and TDP2 genomic regions were additionally excluded.

**Mouse pDAIP-seq processing**

Mouse testis pDAIP-seq reads were processed using the same adapter trimming and fixed 10-nt read-2 trimming strategy used for standard pDAIP-seq and aligned to the mm10 reference genome. PCR duplicates were removed, and alignments with MAPQ ≥10 were retained. Signal tracks were normalized by RPKM. The effective mm10 genome size used for coverage normalization was 2,407,883,243 bp. Peaks were called with the same MACS3 paired-end settings used for human pDAIP-seq and filtered against the corresponding mouse genomic exclusion set, including the TDP1 and TDP2 loci.

**Nucleotide-resolution pDAIP-seq processing and cleavage-site identification**

For nu-pDAIP-seq, reads were adapter- and quality-trimmed with Trim Galore using a quality threshold of 20, stringency 10, a minimum length of 20 nt, and a maximum of 10 ambiguous bases. No fixed 5′ trimming of read 2 was performed because read 2 contains a variable-length Adaptase-derived terminal sequence. Reads were aligned locally to hg38 with Bowtie2, allowing variable terminal sequence to be soft-clipped. PCR duplicates were marked with Picard MarkDuplicates using REMOVE_DUPLICATES=false. Duplicate-marked reads were retained in the primary endpoint analysis, and a sensitivity analysis excluding reads carrying the duplicate flag was performed separately.

Read 2 alignments were selected using SAM flag 128. The first aligned genomic nucleotide of read 2 was extracted at 1-bp resolution using bamCoverage --Offset 1 without read extension. Based on the validated library orientation, read-2 forward alignments corresponded to Crick-strand cleavage products and reverse alignments to Watson-strand cleavage products. Exact cleavage sites were called directly from unsmoothed, unnormalized 1-bp endpoint counts; sites supported by at least two endpoint reads were retained, and sites overlapping the hg38 exclusion set were removed. CPM-normalized tracks smoothed over 11 bp were generated only for visualization.

For cleavage-site sequence analysis, strand-oriented 31-nt genomic sequences comprising positions −16 to −1 and +1 to +15 relative to the cleavage bond were extracted. Minus-strand sequences were reverse-complemented so that all sites were represented in the same cleavage-oriented direction, and only exact 31-nt windows were retained. Sequence logos were generated with ggseqlogo in information-content mode, with the cleavage bond defined between positions −1 and +1.

**Genome-wide signal comparisons and correlations**

Genome-wide comparisons between experimental conditions were generated from spike-in-normalized bigWig tracks using deepTools multiBigwigSummary. Signal was summarized in 10-kb autosomal bins, with chromosomes X, Y, and M excluded and the relevant genomic exclusion regions removed. For condition-to-condition comparisons, bins with normalized signal ≤0.1 in either condition were excluded, values were transformed as log2(signal + 1), and Pearson correlation coefficients were calculated across retained bins. Spearman rank correlations were used for cross-dataset and cross-protein comparisons where indicated in the figures.

**Consensus regions and differential analysis**

For differential analyses, a common region universe was generated by combining TOP1cc peaks from the relevant experimental conditions and merging overlapping intervals with GenomicRanges::reduce(). The resulting union regions were fixed before quantification, and all samples within a comparison were quantified over the same regions using deepTools multiBamSummary and MAPQ-filtered, duplicate-removed BAM files.

Per-sample spike-in read counts obtained with Spiker were appended to each genomic count matrix as a synthetic spike-in control row. Differential analysis was performed with DESeq2 v1.50.1 using a ~ condition design. Size factors were estimated exclusively from the spike-in control row using estimateSizeFactors() with the spike-in row specified as the control feature. The synthetic spike-in row was removed from downstream genomic results. P values were obtained with the DESeq2 Wald test and adjusted for multiple testing by the Benjamini-Hochberg procedure with independent filtering enabled. Log2 fold-change estimates were shrunk using apeglm v1.32.0.

HCT116 triptolide/flavopiridol samples and DLD1 POLR2A-degron samples were modeled separately because they represent distinct cell-line and experimental backgrounds. Cross-system comparisons were made using within-experiment log2 fold changes rather than absolute normalized counts. For the DNA-melted versus DNA-unmelted comparison in Fig. 1, differential regions were defined by |log2 fold change| > 1 and $q < 0.05$.

**Gene-centered profiles and heatmaps**

Gene-centered signal matrices were generated with deepTools computeMatrix scale-regions. For most human TOP1-centered analyses, NCBI RefSeq Select annotation was used with protein-coding canonical genes on autosomes; genes shorter than 1 kb were excluded where indicated. Transcription units were scaled to 6 kb with 3-kb upstream and downstream flanking regions, and signal was summarized in 50-bp bins using the mean signal within each bin. For expression-stratified analyses, genes were classified as high (TPM > 45.5), medium (12.5 < TPM ≤ 45.5), low (0.5 < TPM ≤ 12.5), or silent (TPM ≤ 0.5). Spike-in-normalized tracks were used for cross-condition comparisons, whereas RPKM-normalized tracks were used where RPKM is explicitly indicated.

For cross-topoisomerase analyses in Fig. 7, transcription units were defined using the human NCBI RefSeq Select annotation. For mouse Fig. 7 analyses, transcription units were defined using the mm10 NCBI RefSeq Select annotation. In both species, gene bodies were scaled to 6 kb with 3-kb upstream and downstream flanks and 50-bp bins.

**Cross-topoisomerase analyses**

Genome-wide RPKM-normalized pDAIP-seq signal for TOP1cc, TOP2Acc, TOP2Bcc, TOP3Acc, and TOP3Bcc was summarized in common 10-kb autosomal bins after exclusion of chromosomes X, Y, and M and the hg38 genomic exclusion set. Pairwise associations among the five cleavage-complex profiles were calculated using Spearman rank correlation with pairwise-complete observations. The correlation matrix was ordered for visualization using optimal leaf ordering based on a distance of $1 - \rho$; this ordering was used only for display. A common union of TOP1cc, TOP2Acc, TOP2Bcc, TOP3Acc, and TOP3Bcc regions was used for peak-centered analyses, with RPKM-normalized signal summarized across ±3 kb from region centers in 50-bp bins.

For mouse testis, pairwise correlations among SPO11cc, TOP1cc, TOP2Acc, and TOP2Bcc were calculated from RPKM-normalized signal in 10-kb autosomal bins after exclusion of chromosomes X, Y, and M and the mouse genomic exclusion set. A common union of mouse cleavage-complex regions was used for peak-centered analyses over ±3 kb in 50-bp bins.

##### Genomic-state classification

Genomic states were learned with ChromHMM using binary genomic tracks for ATAC-seq, H3K4me3, H3K27ac, H3K4me1, H3K36me3, H3K9me3, H3K27me3, POLR2A, TOP1, and TOP1cc. Input peak tracks were binarized with ChromHMM BinarizeBed using peak-based input, and a 15-state model was learned on hg38 with LearnModel. States were reordered for presentation according to their emission patterns and genomic annotations, and biological state labels were assigned after inspection rather than automatically. Enrichment of repeat classes and repeat families within each state was calculated with ChromHMM OverlapEnrichment using hg38 RepeatMasker annotations.

##### Repeat-subfamily enrichment

RepeatMasker annotations were used to quantify enrichment of individual repeat subfamilies in TOP1cc-associated regions. Focal TOP1cc regions were compared with one-to-one matched control regions. Candidate control regions were generated from shuffled nonpeak regions, restricted to standard chromosomes, filtered against genomic exclusion regions, required not to overlap observed TOP1cc peaks, and required to have mean mappability ≥0.6. Matching was performed with nullranges::matchRanges using GC content, CpG density, region length, and mappability, with random seed 20202(*12*).

A repeat feature was considered overlapping when the intersection covered at least 1 bp and at least 25% of the focal region, and each focal or control region was counted at most once per feature. Enrichment was expressed as the log2 odds ratio with a Haldane correction of 0.5. The primary paired test was a two-sided exact binomial test of discordant pairs, with Fisher's exact test used as a sensitivity analysis. Benjamini-Hochberg correction was performed separately within the corresponding assay, processing, and annotation groups. High-confidence repeat associations required at least 600 total focal-plus-control observations, at least five observations in each group, FDR < 0.05, and |log2 odds ratio| ≥ 2.

##### LINE-1 analysis

Human L1 loci from L1Base2 were intersected with hg38 RepeatMasker annotations, and each L1Base locus was assigned to the RepeatMasker L1 subfamily contributing the greatest nonredundant overlap. Overlapping RepeatMasker fragments on the same strand were reconstructed within each L1Base insertion. For positional analyses, elements were retained when at least 5 kb of L1 sequence was covered, RepeatMasker fragments covered at least 80% of the reconstructed span, and the reconstructed element consisted of a single continuous fragment on the same strand as the L1Base annotation. Sex-chromosome loci and duplicate reconstructed loci were excluded. Retained elements were grouped by L1 evolutionary subfamily and analyzed in strand-oriented coordinates. Fig. 4 displays L1PA4, L1PA5, L1PA6, L1PA7, and older L1M elements.

L1 profiles were generated with deepTools computeMatrix scale-regions, scaling each L1 body to 6 kb and including 3-kb flanking regions on either side in 50-bp bins. TOP1cc signal was represented by RPKM-normalized tracks.

##### CA–GT sequence-skew analysis

Genome-wide CA–GT skew was defined as $[(A + C) - (G + T)]/(A + C + G + T)$. CA–GT skew was calculated from the hg38 reference sequence using a 200-bp sliding window with a 10-bp step. Non-ACGT bases were excluded from the denominator, and windows containing no canonical A/C/G/T bases were omitted. Both signed and absolute skew tracks were generated; analyses in Fig. 4 used the absolute CA–GT skew.

For the TOP1cc-centered analysis in Fig. 4E, observed TOP1cc regions were compared with positionally shuffled regions. TOP1cc signal and absolute CA–GT skew were summarized around region centers over ±3 kb in 50-bp bins.

##### TOP1cc sequence-motif analysis

Genomic sequences from the top 6,000 endogenous TOP1cc pDAIP-seq regions passing $q < 0.05$ were used as the positive sequence set, and sequences from shuffled TOP1cc regions generated with random seed 60 were used as the negative set. Motif enrichment was analyzed with STREME v5.5.9 in DNA mode using the differential-enrichment objective (--objfun de), random seed 42, motif widths of 6–20 bp, a maximum of 15 motifs, E-value-based significance testing with a threshold of 0.05, and center alignment. STREME motifs were ranked by ascending E value. The third-ranked motif was selected for display and visualized as a sequence logo using ggseqlogo in information-content mode.

##### vipDAIP-seq correlation analysis

Cellular pDAIP-seq and vipDAIP-seq signal was compared within the three amplified genomic intervals shown in Fig. 5. Signal was summarized in 1-kb genomic bins, and Pearson correlation coefficients were calculated between cellular and in vitro TOP1cc signal.

##### Copy-number-matched cross-cell-line analysis

Precomputed segment-level copy-number calls for HCT116 (SIDM00783) and MCF-7 (SIDM00148) were obtained from the Sanger Cell Model Passports/DepMap. Copy-number segments were assigned to prefiltered 10-kb genomic bins using the bin midpoint. Bins lacking a copy-number category in either cell line were excluded. A bin was classified as copy-number matched when the assigned PureCN cn_category was identical between HCT116 and MCF-7. For each autosome, Spearman correlations between HCT116 and MCF-7 TOP1cc signal were calculated using all retained bins and separately using only copy-number-matched bins.

##### Correlation with DNA-end and repair-associated datasets

For comparison of TOP1cc, nucleotide-resolution TOP1cc, TDP1, PARP1, KU70, AP-seq, and GLOE-seq, signal was summarized over a common set of 10-kb genomic windows. Pairwise Spearman correlations were calculated using pairwise-complete observations.

##### Normal-tissue RNA expression and protein abundance

GTEx Analysis V11 median tissue TPM values were used for TOP1, TOP2A, TOP2B, TOP3A, TOP3B, PARP1, XRCC6, and TDP1. Where multiple GTEx subtissues mapped to the same anatomical group, values were summarized by the median. RNA expression was displayed as log2(median TPM + 1), with each gene scaled independently across tissues.

Normal-tissue immunohistochemistry data were obtained from the Human Protein Atlas. Staining categories were converted to ordinal values of 0, 1, 2, and 3 corresponding to not detected, low, medium, and high staining, respectively. When multiple cell types or source tissues mapped to a single anatomical group, the maximum staining category was retained. IHC data were not available for TOP3B.

Mass-spectrometry protein-abundance data were obtained from the Human Protein Atlas. Replicate measurements for each protein and tissue were summarized by the median, and multiple source tissues mapping to the same anatomical group were further summarized by the median. Protein abundance was displayed as log10(intensity + 1). Missing measurements were retained as missing rather than assigned zero. Anatomical maps used a common scale for female and male maps within each gene or protein. Female maps display tissues shared between sexes together with female-specific tissues; male maps display male-specific tissues.

**Software**

Sequence processing and genomic analyses were performed using Trim Galore v0.6.10, Cutadapt v4.9, Bowtie2 v2.5.4, SAMtools v1.16.1, Picard v2.27.5, deepTools v3.5.5, BEDTools v2.31.1, MACS3 v3.0.2, pyGenomeTracks v3.9, and Spiker v1.0.3. Motif enrichment was performed using STREME v5.5.9. Statistical analyses were performed in R v4.2.1. Differential analyses used DESeq2 v1.50.1 and apeglm v1.32.0. Data visualization used ggplot2 v4.0.3, and sequence logos were generated using ggseqlogo v0.2.2.

#### Supplemental Tables

##### Table S1. Antibodies used in this study

| Target | Supplier | Catalog no. | Application |
|---|---|---|---|
| TOP1, antibody 1 | Novus Biologicals | NBP1-30482 | pDAIP-seq; nu-pDAIP-seq |
| TOP1, antibody 2 | Abcam | ab109374 | pDAIP-seq; ChIP-seq |
| TOP1, antibody 3 | MilliporeSigma | MABE1084 | nu-pDAIP-seq |
| TOP2A | Abcam | ab52934 | pDAIP-seq; immunoblot |
| TOP2B | Abcam | ab72334 | pDAIP-seq |
| TOP2B | BD Biosciences | 611492 | pDAIP-seq |
| TOP3A | Proteintech | 14525-1-AP | pDAIP-seq |
| TOP3B | Abcam | ab183520 | pDAIP-seq |
| TDP1 | Cell Signaling Technology | 59710S | pDAIP-seq |
| PARP1 | Cell Signaling Technology | 9532S | pDAIP-seq |
| KU70/XRCC6 | Cell Signaling Technology | 4588S | pDAIP-seq |
| SPO11 | MilliporeSigma | MABE1167 | mouse-testis pDAIP-seq |
| POLR2A | Abcam | ab5408 | immunoblot |
| GFP | Novus Biologicals | NB600-308 | immunoblot |
| ACTB/β-Actin | Santa Cruz Biotechnology | sc-47778 | immunoblot |
| Anti-RNA–DNA hybrid (S9.6) | Kerafast | ENH001 | DRIP-Rx |
| Rabbit IgG | ABclonal | AC005 | IP control |
| Mouse IgG | Santa Cruz Biotechnology | sc-2025 | IP control |
| ChIP Spike-in Antibody | Active Motif | 61686 | pDAIP-seq; DRIP-Rx normalization |
| Goat Anti-Rabbit IgG H&L (HRP) | Abcam | ab6721 | immunoblot |
| Goat Anti-Mouse IgG H&L (HRP) | Abcam | ab6789 | immunoblot |

#### Table S2. Key reagents and resources

| Reagent/resource | Supplier | Catalog no. |
|---|---|---|
| DNAzol Reagent | Thermo Fisher Scientific | 10503027 |
| RNase A, DNase/protease free | Thermo Fisher Scientific | EN0531 |
| Proteinase K | Thermo Fisher Scientific | AM2544 |
| Pierce Protein A/G Magnetic Beads | Thermo Fisher Scientific | 88803 |
| Dynabeads MyOne Streptavidin C1 | Thermo Fisher Scientific | 65001 |
| Qubit dsDNA Broad Range Assay | Thermo Fisher Scientific | Q33266 |
| SPRIselect | Beckman Coulter | B23318 |
| VAHTS ssDNA Library Prep Kit for Illumina | Vazyme | ND620-02 |
| xGen ssDNA and Low-Input DNA Prep Kit | IDT | 10009817 |
| Human topoisomerase I | TopoGEN | TG2005H-RC2 |
| Human TDP1 | TopoGEN | TG2004H-2 |
| Human TDP2 | TopoGEN | TG2003H-2 |
| ChIP Spike-in Chromatin | Active Motif | 53083 |
| N3-kethoxal | ApexBio | A8793 |
| DBCO-PEG4-Biotin | Jena Bioscience | CLK-A105P4-10 |
| RNase T1 | Thermo Fisher Scientific | EN0541 |
| ShortCut RNase III | New England Biolabs | M0245S |
| RNase H | New England Biolabs | M0297S |
| Micrococcal nuclease | New England Biolabs | M0247 |
| Benzonase Nuclease HC | Sigma-Aldrich | 71205-3 |
| LongAmp Taq 2X Master Mix | New England Biolabs | M0287S |
| KOD One PCR Master Mix | TOYOBO | KMM-101NV |
| Flavopiridol | Santa Cruz Biotechnology | sc-202157A |
| Triptolide | Sigma-Aldrich | T3652 |
| Camptothecin | Sigma-Aldrich | 208925 |
| Doxycycline hyclate | Sigma-Aldrich | D9891-10G |
| 1-Naphthaleneacetic acid (NAA) | Santa Cruz Biotechnology | sc-253943 |
| Auxinole | MedChemExpress | HY-111444 |
| Fetal bovine serum | Thermo Fisher Scientific | 26140079 |
| Penicillin-Streptomycin | Sigma-Aldrich | P4333 |
| McCoy's 5A Modified Medium | Thermo Fisher Scientific | 16600108 |
| Dulbecco's Modified Eagle Medium (DMEM) | Thermo Fisher Scientific | 11965126 |
| 10× RIPA buffer | Sigma-Aldrich | 20-188 |
| DC™ Protein Assay Kit II | Bio-Rad | 5000112 |

#### Table S3. vipDAIP-seq primer sequences

| Locus | Primer | Sequence (5′→3′) |
|---|---|---|
| ABHD10 | abhd10_locus.f | AAGTGGAGGACGGCTTCATTTAA |
| ABHD10 | abhd10_locus.r | GTCTGAAAGATGTCGGTGGTTCT |
| HIVEP3 | hivep3_locus.f | TGGCCTGATCTCTTGTCAGTTTCT |
| HIVEP3 | hivep3_locus.r | CCACCTGGCCTTCTGAGAT |
| MASP2 | masp2_locus.f | TGGCCAAGTTGCCCATTTTAATCA |
| MASP2 | masp2_locus.r | CTCCACCCGGCATTCAGACT |

### Supplemental Figures

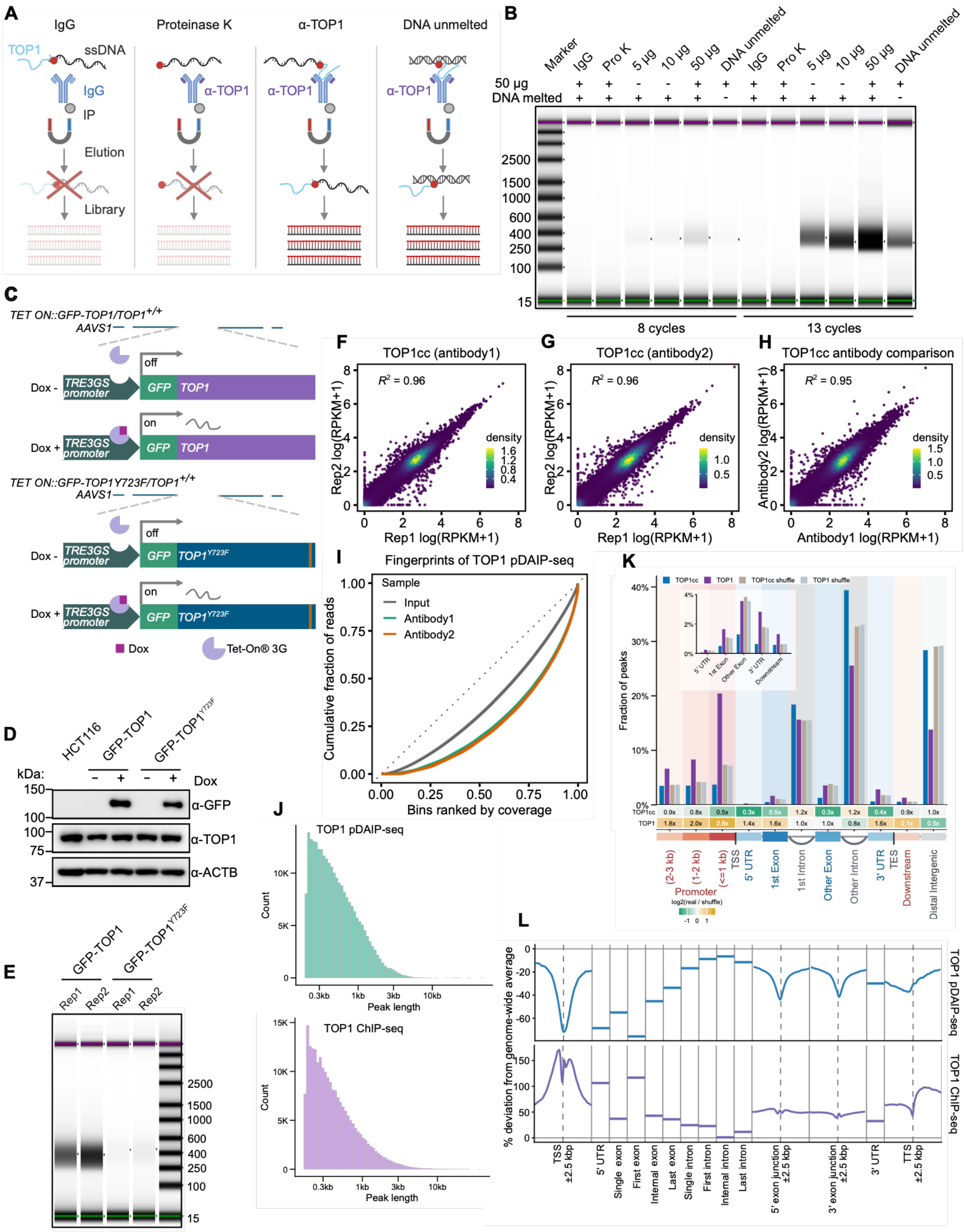

**Fig. S1. Validation and controls for pDAIP-seq to capture endogenous TOP1cc**

**(A)** Schematic of pDAIP-seq specificity controls, including IgG IP, proteinase K pretreatment before IP to remove TOP1–DNA adducts, and omission of DNA melting. **(B)** TapeStation fragment-size profiles of libraries generated under the indicated conditions. **(C)** Schematics of doxycycline (Dox)-inducible GFP-TOP1 and catalytically inactive GFP-TOP1$^{Y723F}$ constructs integrated at the AAVS1 locus in HCT116 cells. **(D)** Immunoblot confirming Dox-dependent expression of GFP-TOP1 and GFP-TOP1$^{Y723F}$. **(E)** TapeStation fragment-size profiles from two replicates of GFP-TOP1 and GFP-TOP1$^{Y723F}$ pDAIP-seq samples. **(F and G)** Genome-wide reproducibility of TOP1cc pDAIP-seq signal between biological replicates using two independent TOP1 antibodies. **(H)** Genome-wide comparison of TOP1cc signal obtained with the two TOP1 antibodies. **(I)** Fingerprint plots showing the cumulative fraction of reads across bins ranked by coverage for input and TOP1cc pDAIP-seq generated with the two antibodies. **(J)** Peak-length distributions for TOP1 pDAIP-seq and TOP1 ChIP-seq. **(K)** Genomic annotation of TOP1cc pDAIP-seq and TOP1 ChIP-seq peaks compared with matched shuffled regions; enrichment or depletion for each annotation category is shown as log2(real/shuffle), with low-frequency categories enlarged in the inset. **(L)** Relative distribution of TOP1 pDAIP-seq and TOP1 ChIP-seq signal across transcription unit features and flanking regions, shown as percent deviation from the genome-wide average.

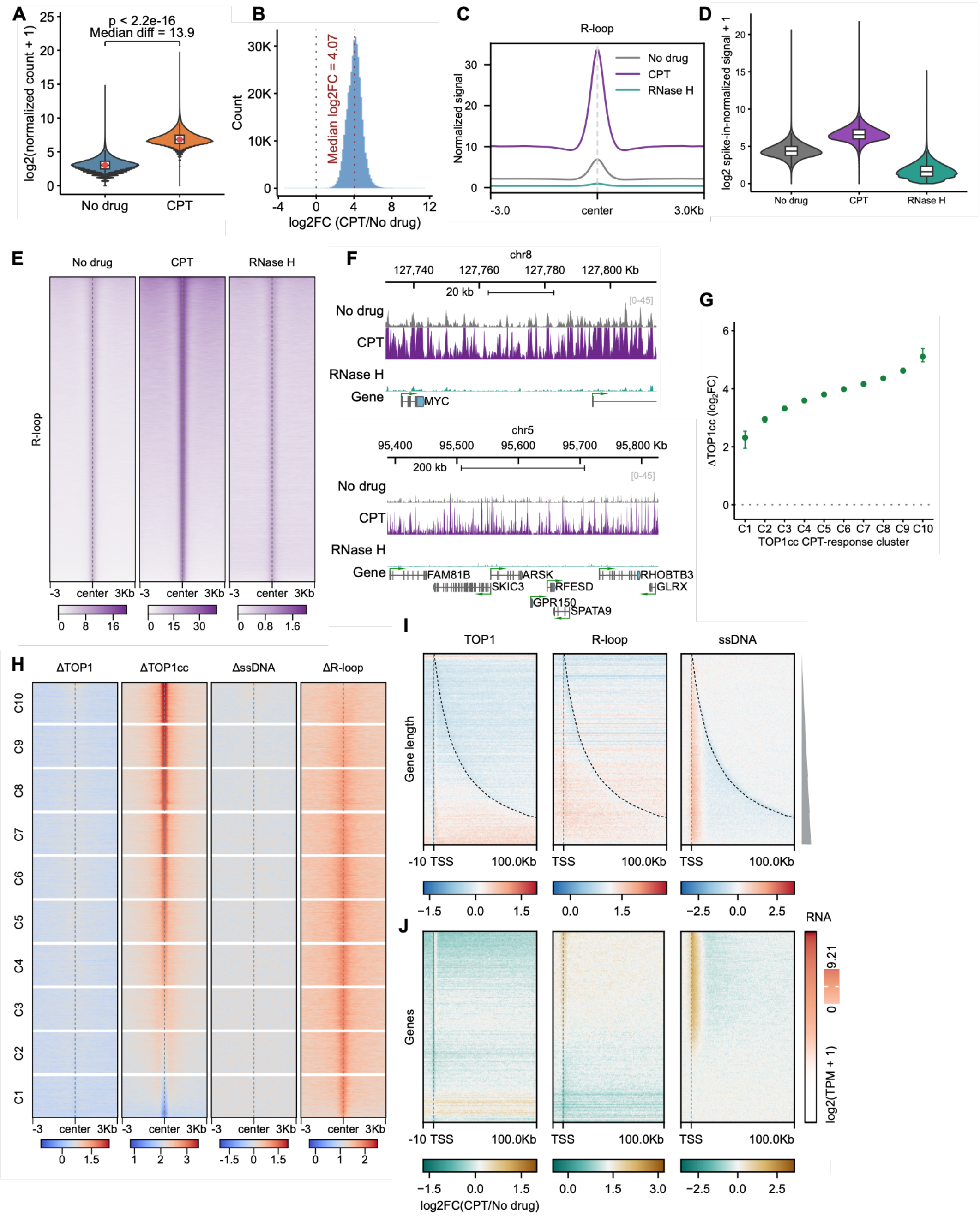

A
p < 2.2e-16
Median diff = 13.9
log2(normalized count + 1)
No drug
CPT
B
Median log2FC = 4.07
Count
log2FC (CPT/No drug)
C
R-loop
Normalized signal
No drug
CPT
RNase H
-3.0
center
3.0Kb
D
log2 spike-in-normalized signal + 1
No drug
CPT
RNase H
E
No drug
CPT
RNase H
R-loop
F
chr8
20 kb
No drug
CPT
RNase H
Gene
MYC
chr5
200 kb
FAM81B
ARSK
RHOBTB3
SKIC3
RFESD
GLRX
GPR150
SPATA9
G
ΔTOP1cc (log2FC)
TOP1cc CPT-response cluster
H
ΔTOP1
ΔTOP1cc
ΔssDNA
ΔR-loop
I
TOP1
R-loop
ssDNA
Gene length
J
Genes
RNA
log2(TPM + 1)
log2FC(CPT/No drug)

**Fig. S2. CPT-induced genome-wide alterations of TOP1cc, DRIP-Rx signal, and ssDNA.**
**(A)** Genome-wide distribution of normalized TOP1cc pDAIP-seq counts in no-drug and camptothecin (CPT)-treated cells. **(B)** Distribution of TOP1cc log2(CPT/no drug) fold changes. **(C)** Metaprofiles of DRIP-Rx signal centered on R-loop regions in no-drug, CPT-treated, and RNase H-treated samples. **(D)** Spike-in-normalized DRIP-Rx signal across the indicated conditions. **(E)** DRIP-Rx heatmaps centered on R-loop regions. **(F)** Representative genome-browser views of DRIP-Rx signal at the MYC locus on chromosome 8 and a locus on chromosome 5. **(G)** Mean TOP1cc change across ten CPT-response clusters. **(H)** Heatmaps of CPT-induced changes in TOP1, TOP1cc, ssDNA, and DRIP-Rx signal across the ten CPT-response clusters. **(I-J)** Heatmaps of CPT-induced changes in TOP1, DRIP-Rx signal, and ssDNA aligned to transcription start sites (TSSs) and extending 100 kb downstream, with genes ordered by gene length **(I)** or RNA abundance **(J)**.

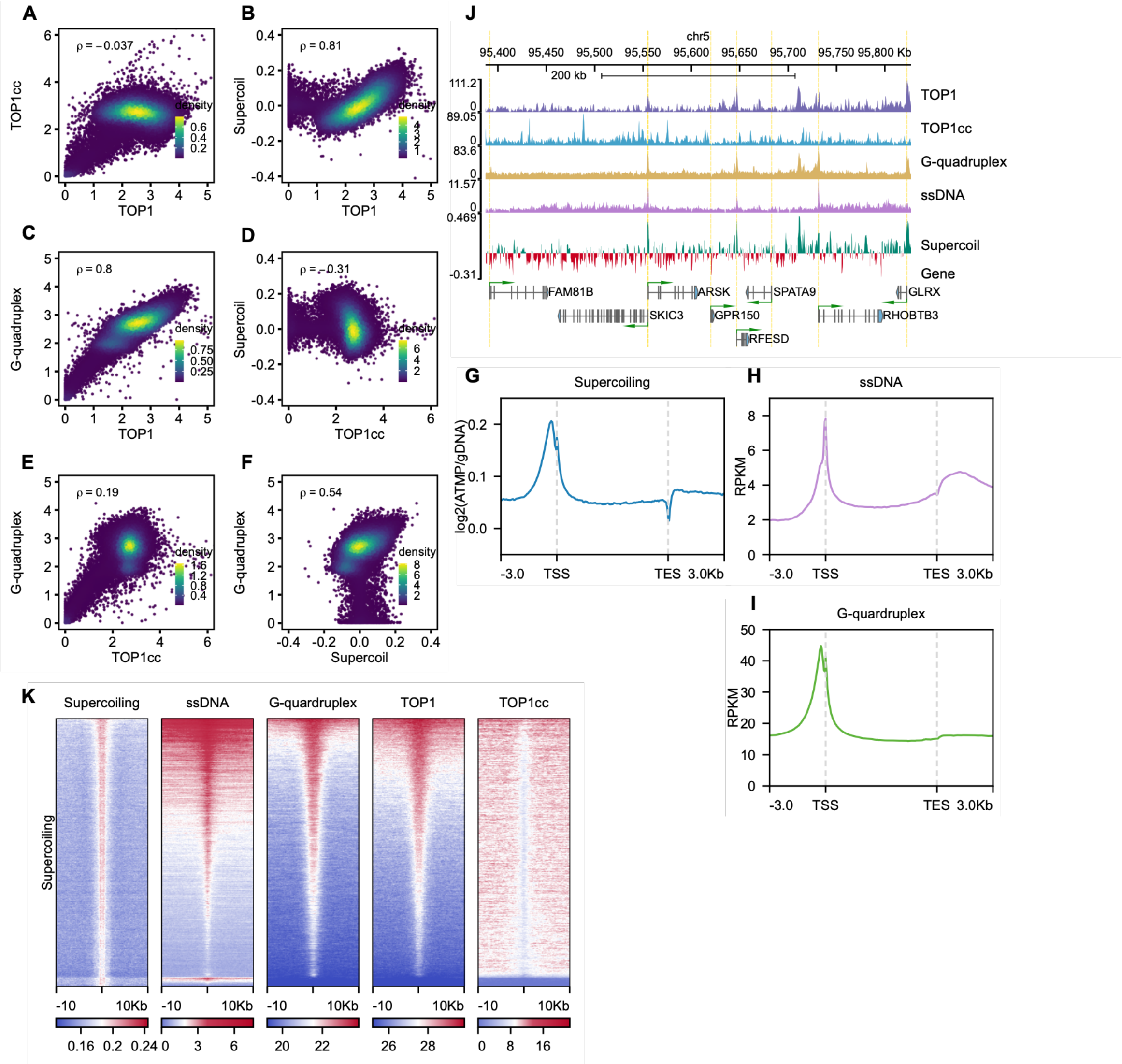

**Fig. S3. Relationships between TOP1 binding, TOP1cc, DNA supercoiling, ssDNA, and G-quadruplexes**

**(A–F)** Genome-wide pairwise comparisons of TOP1, TOP1cc, supercoiling, and G-quadruplex signals; Spearman correlation coefficients (ρ) are indicated. **(A)** TOP1 versus TOP1cc. **(B)** TOP1 versus supercoiling. **(C)** TOP1 versus G-quadruplexes. **(D)** TOP1cc versus supercoiling. **(E)** TOP1cc versus G-quadruplexes. **(F)** Supercoiling versus G-quadruplexes. **(G–I)** Metagene profiles of supercoiling **(G)**, ssDNA **(H)**, and G-quadruplex **(I)** signals across transcription units and 3-kb flanking regions. **(J)** Representative genome-browser view on chromosome 5 showing TOP1, TOP1cc, G-quadruplex, ssDNA, supercoiling, and gene tracks. **(K)** Heatmaps of supercoiling, ssDNA, G-quadruplex, TOP1, and TOP1cc signals centered on supercoiling-associated regions with 10-kb flanking regions.

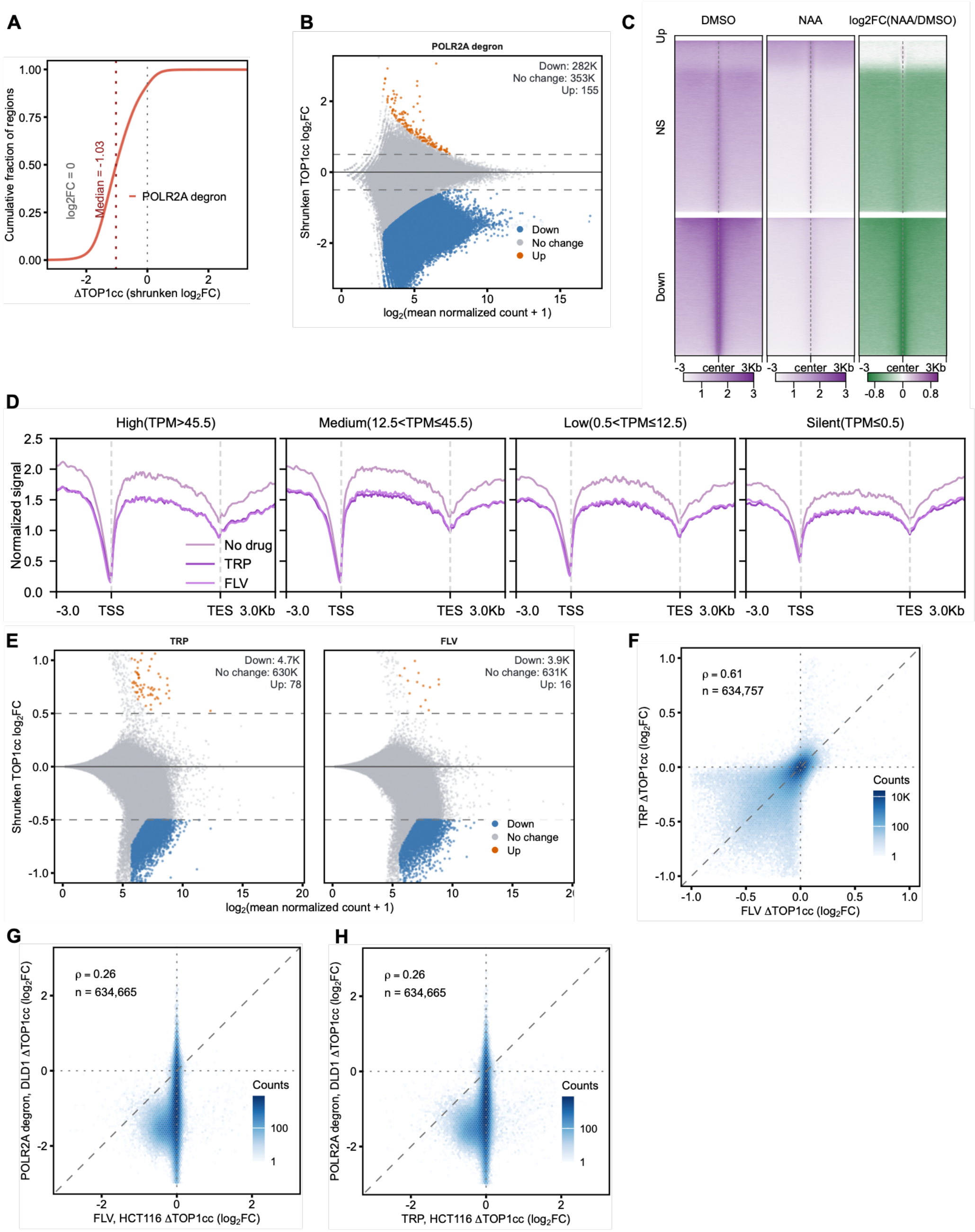
A
B
C
D
E
F
G
H
POLR2A degron
Down: 282K
No change: 353K
Up: 155
Down
No change
Up
DMSO
NAA
log2FC(NAA/DMSO)
NS
High(TPM>45.5)
Medium(12.5<TPM≤45.5)
Low(0.5<TPM≤12.5)
Silent(TPM≤0.5)
No drug
TRP
FLV
TRP
FLV
Down: 4.7K
No change: 630K
Up: 78
Down: 3.9K
No change: 631K
Up: 16
ρ = 0.61
n = 634,757
ρ = 0.26
n = 634,665
ρ = 0.26
n = 634,665
Counts

**Fig. S4. POLR2A degradation and transcription inhibition reduce global TOP1cc abundance**

**(A)** Cumulative distribution of shrunken TOP1cc log[1]2 fold changes after POLR2A degradation. **(B)** MA plot of TOP1cc changes after POLR2A degradation. **(C)** Heatmaps of TOP1cc signal in DMSO- and NAA-treated cells and the corresponding log2(NAA/DMSO) fold change. **(D)** Metagene profiles of TOP1cc signal after treatment with triptolide (TRP), flavopiridol (FLV), or no drug control, stratified by RNA abundance. **(E)** MA plots of TOP1cc changes after TRP or FLV treatment. **(F)** Genome-wide comparison of TOP1cc changes induced by TRP and FLV in HCT116 cells. **(G and H)** Genome-wide comparisons of TOP1cc changes after POLR2A degradation in DLD1 cells with those induced by FLV **(G)** or TRP **(H)** in HCT116 cells.

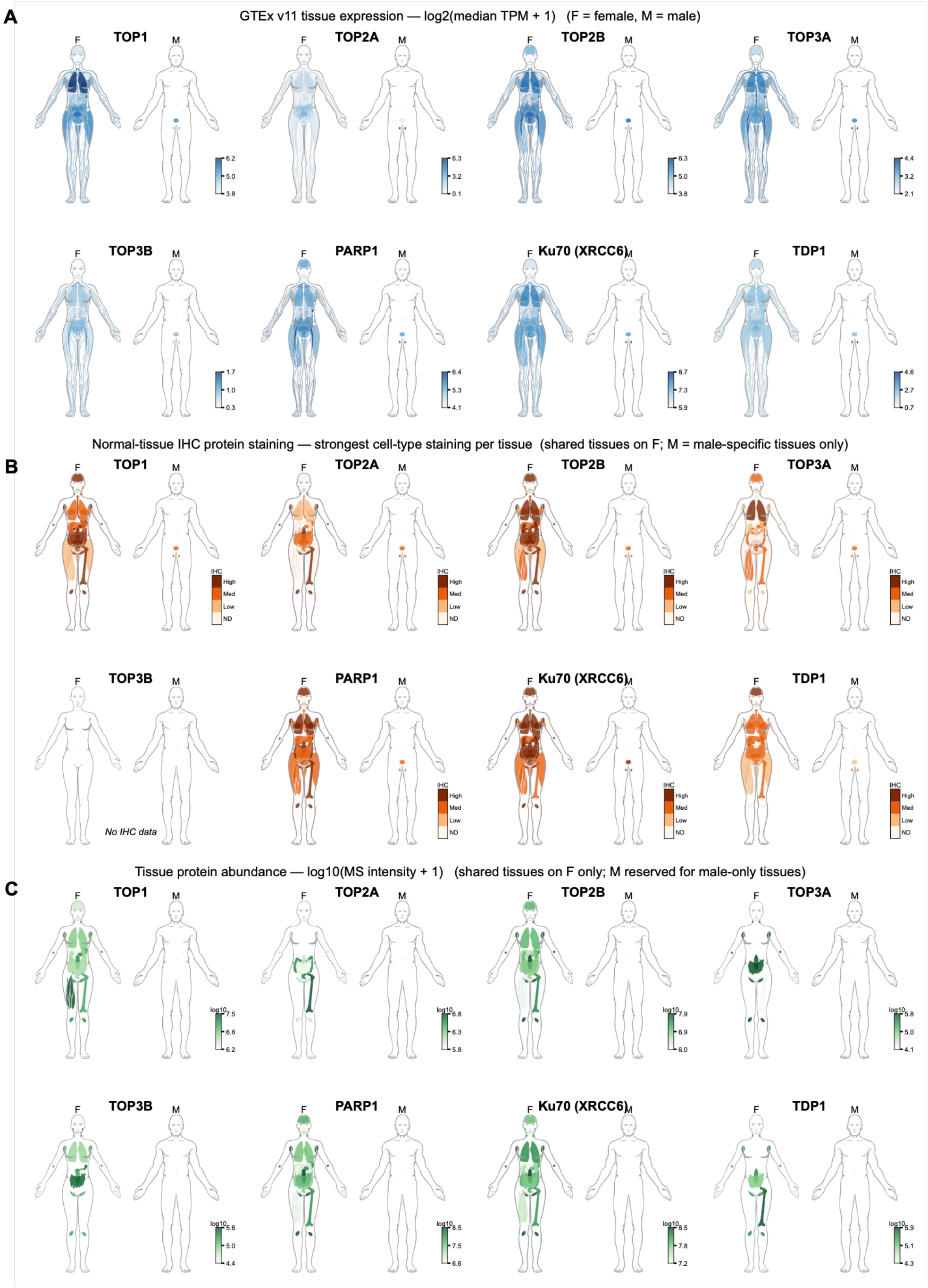
A
GTEx v11 tissue expression — log2(median TPM + 1)   (F = female, M = male)
TOP1
TOP2A
TOP2B
TOP3A
TOP3B
PARP1
Ku70 (XRCC6)
TDP1
B
Normal-tissue IHC protein staining — strongest cell-type staining per tissue  (shared tissues on F; M = male-specific tissues only)
IHC
High
Med
Low
ND
No IHC data
C
Tissue protein abundance — log10(MS intensity + 1)   (shared tissues on F only; M reserved for male-only tissues)
log10

**Fig. S5. Tissue expression patterns and protein abundance of topoisomerases and DNA repair factors**

(A–C) Anatomical maps show GTEx v11 median tissue RNA expression (A), strongest cell-type immunohistochemical (IHC) staining in normal tissues (B), and mass-spectrometry (MS) protein abundance (C) for TOP1, TOP2A, TOP2B, TOP3A, TOP3B, PARP1, KU70 (XRCC6), and TDP1. Female maps show tissues shared between sexes; male maps show male-specific tissues. IHC data were not available for TOP3B.
